# Premeiotic 24-nt phasiRNAs are present in the *Zea* genus and unique in biogenesis mechanism and molecular function

**DOI:** 10.1101/2024.03.29.587306

**Authors:** Junpeng Zhan, Sébastien Bélanger, Scott Lewis, Chong Teng, Madison McGregor, Aleksandra Beric, Michael A. Schon, Michael D. Nodine, Blake C. Meyers

**Affiliations:** National Key Laboratory of Crop Genetic Improvement, Huazhong Agricultural University, Wuhan 430070, China; Hubei Hongshan Laboratory, Wuhan 430070, China; Donald Danforth Plant Science Center, St. Louis, MO 63132, USA; The James Hutton Institute, Dundee, Scotland DD2 5DA, UK; Division of Biology and Biomedical Sciences, Washington University, St. Louis, MO 63130, USA; Division of Plant Science and Technology, University of Missouri, Columbia, MO 65211, USA; Laboratory of Molecular Biology, Wageningen University, Wageningen 6708 PB, the Netherlands

**Keywords:** Maize, teosinte, small RNA, phasiRNA, nanoPARE

## Abstract

Reproductive phasiRNAs are broadly present in angiosperms and play crucial roles in sustaining male fertility. While the premeiotic 21-nt phasiRNAs and meiotic 24-nt phasiRNA pathways have been extensively studied in maize (*Zea mays*) and rice (*Oryza sativa*), a third putative category of reproductive phasiRNAs–named premeiotic 24-nt phasiRNAs–have recently been reported in barley (*Hordeum vulgare*) and wheat (*Triticum aestivum*). To determine whether premeiotic 24-nt phasiRNAs are also present in maize and related species and begin to characterize their biogenesis and function, we performed a comparative transcriptome and degradome analysis of premeiotic and meiotic anthers from five maize inbred lines and three teosinte species/subspecies. Our data indicate that a substantial subset of the 24-nt phasiRNA loci in maize and teosinte are already highly expressed at premeiotic phase. The premeiotic 24-nt phasiRNAs are similar to meiotic 24-nt phasiRNAs in genomic origin and dependence on DCL5 for biogenesis, however, premeiotic 24-nt phasiRNAs are unique in that they are likely (i) not triggered by microRNAs, (ii) not loaded by AGO18 proteins, and (iii) not capable of mediating *cis*-cleavage. In addition, we also observed a group of premeiotic 24-nt phasiRNAs in rice using previously published data.

Together, our results indicate that the premeiotic 24-nt phasiRNAs constitute a unique class of reproductive phasiRNAs and are present more broadly in the grass family (Poaceae) than previously known.

**SIGNIFICANCE:** We previously reported two classes of reproductive phasiRNAs in maize, the premeiotic 21-nt phasiRNAs and the meiotic 24-nt phasiRNAs. Here we report a third class of reproductive phasiRNAs – premeiotic 24-nt phasiRNAs – that are present in the *Zea* genus, including all five maize inbred lines and three teosinte species/subspecies that we examined, plus rice.

We show that in the *Zea* genus the premeiotic 24-nt phasiRNAs are distinct from the meiotic 24-nt phasiRNAs in triggering mechanism, effector protein, and molecular function.

## INTRODUCTION

Two size classes of reproductive, phased, small interfering RNAs (phasiRNAs) – 21 nucleotides (nt) and 24 nt in length – accumulate to high abundance in the anthers of numerous flowering plants. The total abundance of the 21-nt phasiRNAs usually peaks at the premeiotic phase of anther development, whereas the 24-nt phasiRNAs peak during meiosis (1). The premeiotic 21-nt phasiRNAs and meiotic 24-nt phasiRNAs are derived from long noncoding loci and produced via two genetically separable biogenesis pathways. In both pathways, phasiRNA precursors (aka *PHAS* precursors) are generated from *PHAS* loci by RNA polymerase II, cleaved by an Argonaute1 (AGO1) clade protein directed by a microRNA (miRNA), converted to double stranded RNA molecules, and chopped by a Dicer-like (DCL) protein to produce small interfering RNAs (siRNAs) that map to the corresponding genomic loci in a precise head-to-tail arrangement (hence the term “phased siRNA”). The two pathways differ primarily in miRNA triggers and DCL proteins, with miR2118 and DCL4 acting in the 21-nt phasiRNA pathway, and miR2275 and DCL5 acting in the meiotic 24-nt phasiRNA pathway (2). The AGO proteins known to load 21-nt phasiRNAs and 24-nt phasiRNAs include a few pathway-specific members and a few that are common to both pathways. For example, AGO1d loads 21-nt phasiRNAs in rice (3, 4); AGO5 clade proteins load 21-nt phasiRNAs in rice and maize (5, 6); and AGO18b loads both 21-nt and 24-nt phasiRNAs in maize (7).

Reproductive phasiRNAs are crucial for maintaining male fertility. Mutations in a few rice 21-nt phasiRNA loci (aka *21-PHAS* loci) cause temperature / photoperiod-sensitive male sterility (8, 9), and loss of a subset of MIR2118 genes in rice causes male and female sterility (10). In maize, loss-of-function mutants of *Dcl5*, which likely functions specifically in the 24-nt phasiRNA pathway, exhibit temperature-sensitive male sterility (11). In terms of molecular functions, the 21-nt phasiRNAs are known to mediate *cis*-cleavage of their own precursors in rice and maize (12), and *trans*-cleavage of protein-coding mRNAs in rice (13, 14). In maize, 24-nt phasiRNAs are essential for maintaining CHH DNA methylation at their own genomic loci in *cis* (15). However, RNA targets of the 21-nt and 24-nt phasiRNAs in maize, if any, remain largely unknown.

The 21-nt phasiRNAs and meiotic 24-nt phasiRNAs both have been demonstrated to be widely present in angiosperms (16). However, their patterns of conservation differ; except for a handful of eudicot species that have recently been shown to accumulate 21-nt phasiRNAs (17), the majority of eudicots lack 21-nt phasiRNAs, whereas the meiotic 24-nt phasiRNAs are more broadly present in angiosperms. Notably, several eudicot species– including the model plant species *Arabidopsis thaliana*–lack both 21- and 24-nt phasiRNAs (16). We previously reported a group of 24-nt phasiRNAs that peak at the premeiotic phase of anther development in barley and wheat and lack miR2275 target sites (18). However, whether the premeiotic 24-nt phasiRNAs are also present in other grass lineages remains poorly understood, as does their biogenesis mechanisms and functions.

Here, we carried out a comprehensive survey of reproductive phasiRNA pathways in five maize inbred lines and three teosinte species/subspecies. The maize inbred lines were chosen from among the founders of the maize nested association mapping population, including a stiff-stalk (B73), a non-stiff-stock (Oh43), a popcorn (HP301), a sweet corn (Il14H), and a tropical (NC358) variety, each representing a major clade of modern maize inbred lines (19, 20), and the teosinte varieties included two subspecies – *Zea mays* ssp. *parviglumis* (TIL11) and *Zea mays* ssp. *mexicana* (TIL25) – that are known as progenitors of modern maize (21), and *Zea luxurians* (RIL003). We show that in all eight *Zea* varieties, a substantial subset of *24-PHAS* loci are highly expressed at the premeiotic phase of anther development. These premeiotic 24-nt phasiRNAs are distinct from the meiotic 24-nt phasiRNAs and premeiotic 21-nt phasiRNAs in several aspects of biogenesis mechanisms and functions, and thus constitute a unique class of reproductive phasiRNAs.

## RESULTS

### Conservation of phasiRNA pathway genes in *Zea* genus

To identify reproductive *PHAS* loci and phasiRNA pathway genes in the eight *Zea* varieties (Figure 1A), we performed small RNA-seq (sRNA-seq), RNA-seq, and nanoPARE analyses of 2 to 5 developmental stages of anthers in each variety, spanning premeiotic to meiotic phases (Figure 1B). We annotated the HD-ZIP IV, bHLH, RDR, DCL, AGO, SGS3, DRB, SE, SDN1 and HESO1/URT1 family proteins, which are involved in the biogenesis of phasiRNAs and/or miRNAs (2), encoded in all the *Zea* genomes and a few outgroup species, and performed phylogenetic analyses for each family (Figure 1A; Figure S1). The gene copy numbers of each major clade in each phylogenetic tree are similar across the *Zea* genomes, with only a few clades (e.g. the RDR6, DRB4, and HEN1 clades) exhibiting copy number variation (Dataset S1). These data suggest that the phasiRNA pathway genes are largely conserved in the *Zea* genus.

**Figure 1.**
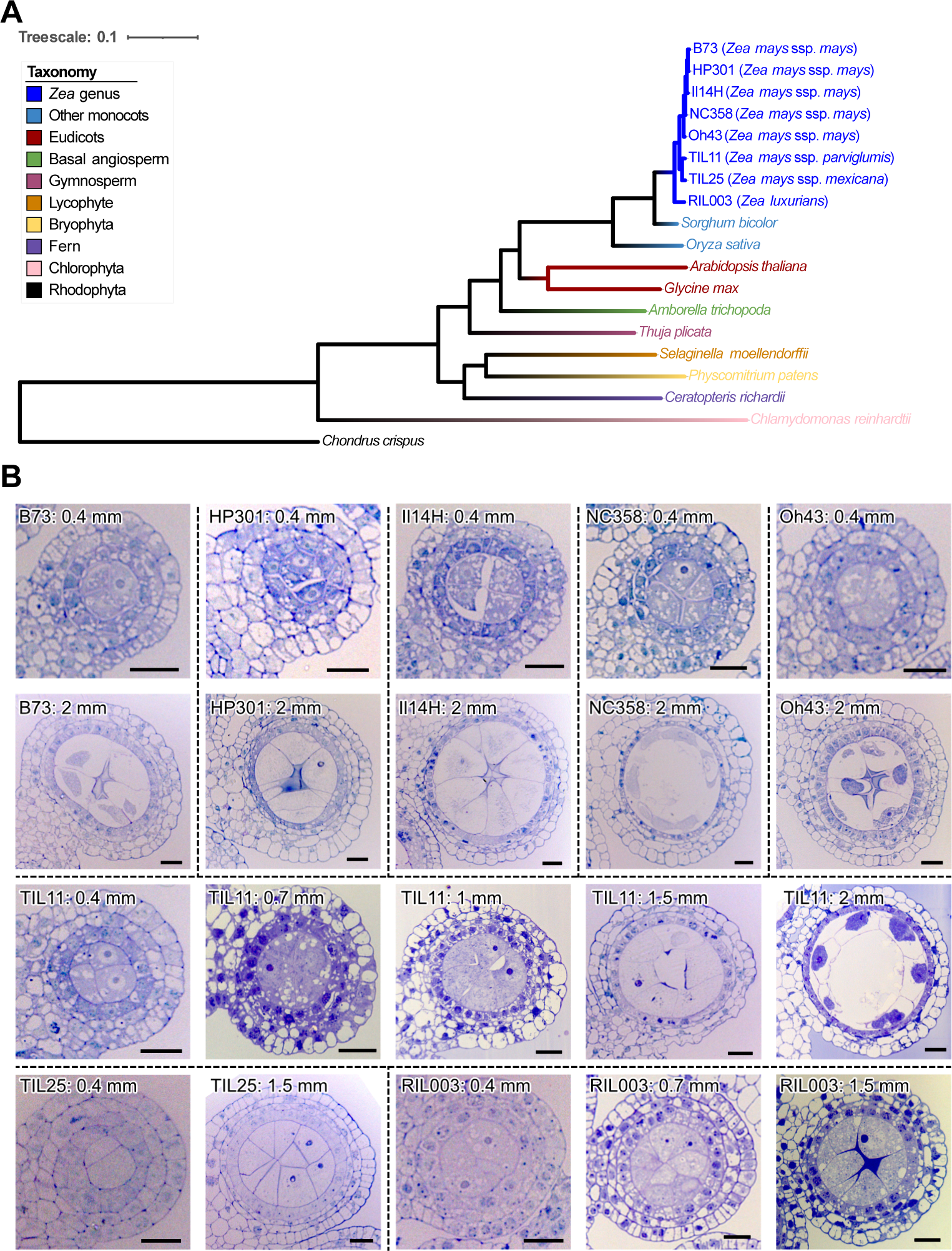
*Zea* varieties and anther developmental stages sampled for the comparative analysis of phasiRNAs. **(A)** Phylogeny of the *Zea* varieties analyzed in this study and outgroup species used in the phylogenetic analysis of gene families. **(B)** Micrographs of cross sections of anthers representing those sampled for transcriptome and degradome analyses. In the maize varieties (B73, HP301, Il14H, NC358, and Oh43) and TIL25, 0.4-mm anthers are premeiotic and 2-mm anthers are meiotic. In TIL11, 0.4 to 0.7-mm anthers are premeiotic, and 1.5- to 2-mm anthers are meiotic. In RIL003, 0.4- and 0.7-mm anthers are premeiotic, and 1.5-mm anthers are meiotic. Scale bars = 20 μm.

### Identification of reproductive *PHAS* loci in *Zea* varieties

Using the sRNA-seq data, we identified reproductive *PHAS* loci and examined their temporal expression patterns during early anther development in the *Zea* varieties. Similar to the W23 *bz2* maize (1), we found that in the five maize inbred lines plus TIL11, 21-nt phasiRNAs were more abundant in premeiotic anthers than in meiotic anthers. However, interestingly, in TIL25 and RIL003, 21-nt phasiRNAs were more abundant at the meiotic phase than the premeiotic phase (Figure S2). These results suggest that the gene regulatory program controlling the temporal accumulation of 21-nt phasiRNAs diverged between TIL11 and the other two teosinte varieties, and the TIL11-like program was retained in modern maize. Remarkably, while it had previously been demonstrated in maize that the 24-nt phasiRNAs are most abundant during meiosis (1), in all the *Zea* varieties we analyzed, we observed a group of *24- PHAS* loci that already accumulated a high abundance of phasiRNAs at the premeiotic phase (Figure 2A; Dataset S2). Hereafter, the *PHAS* loci that produce substantial levels of phasiRNAs (CPM > 20) at the premeiotic stage are referred to as premeiotic *24-PHAS* loci, and the other *24-PHAS* loci are referred to as meiotic *24-PHAS* loci. The two classes of *24- PHAS* loci and the *21-PHAS* loci do not overlap in genomic locations in any of the *Zea* varieties. In TIL11 and the modern maize varieties, phasiRNA abundance of the premeiotic *24-PHAS* loci declined substantially by the meiotic phase, whereas this decrease did not occur or was modest in TIL25 and RIL003 (Figure 2A), suggesting that TIL11 evolved a regulatory mechanism that downregulates the abundance of 24-nt phasiRNAs after the premeiotic phase, and such a mechanism has been maintained in modern maize.

**Figure 2.**
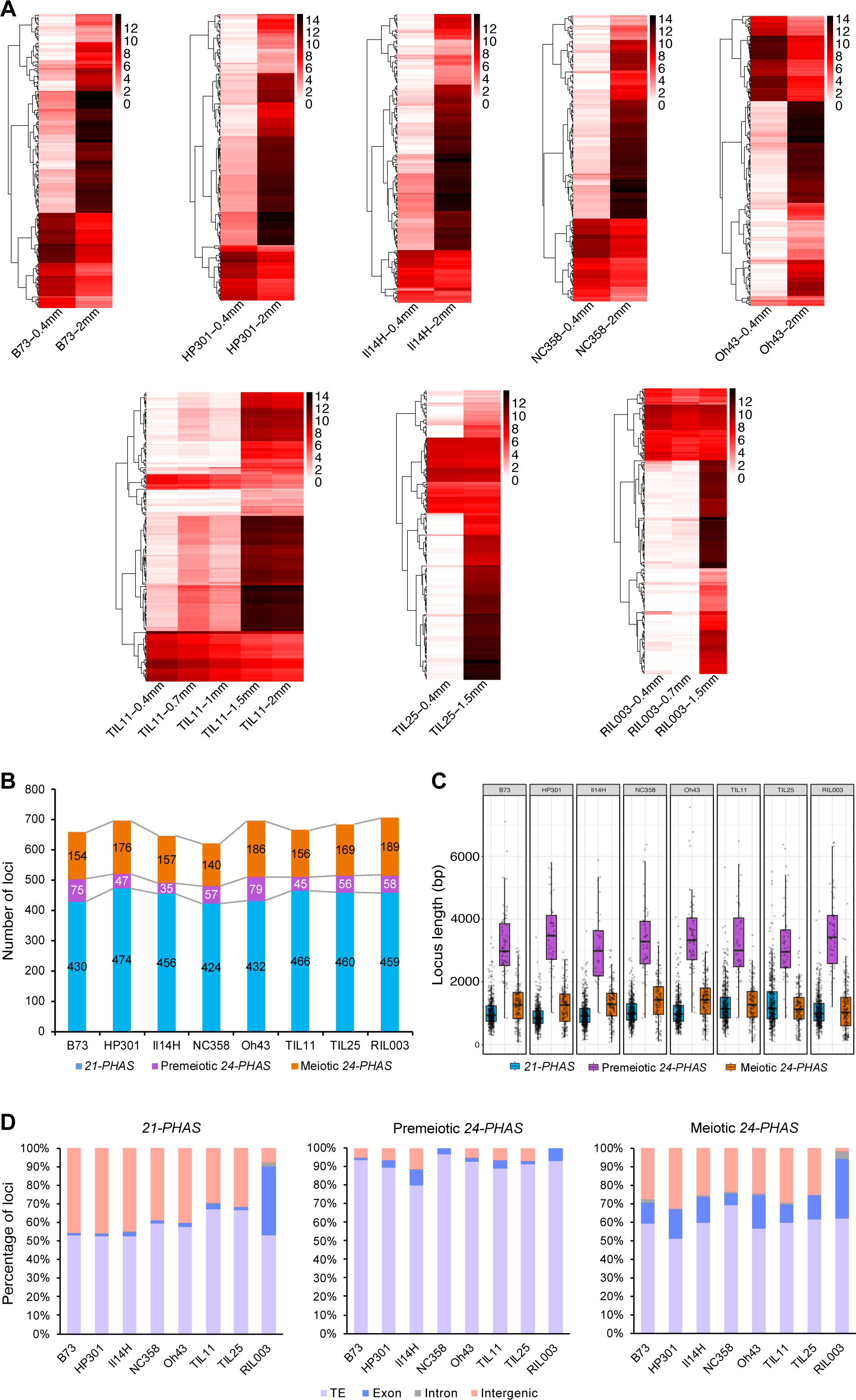
Identification of reproductive *PHAS* loci in the *Zea* varieties. **(A)** Heatmaps of abundance of phasiRNAs derived from individual *24-PHAS* loci. Heatmaps were clustered on Euclidean distance. **(B)** Numbers of *PHAS* loci in each genome. **(C)** Boxplot of *PHAS* loci lengths. **(D)** Percentages of *PHAS* loci overlapping with TEs, exons, introns, or intergenic regions.

The numbers of *21-* and *24-PHAS* loci were similar across the *Zea* varieties and to the previously reported loci numbers in W23 *bz2*, which has 463 *21-PHAS* and 176 *24-PHAS* loci (1) (Figure 2B). Interestingly, in every variety, the premeiotic *24-PHAS* loci were significantly longer than the *21-PHAS* and meiotic *24-PHAS* loci (*P* < 0.05) (Figure 2C; Dataset S3). The distribution of *PHAS* loci across chromosomes was similar among the *Zea* varieties in that (i) in every variety, the *21-PHAS*, premeiotic *24-PHAS*, and meiotic *24-PHAS* were distributed across all chromosomes (Figures S3 and S4); and (ii) in nearly every variety, Chr. 2 had the largest number of *21-PHAS* loci, Chr. 4 had the largest number of premeiotic *24-PHAS* loci, and Chr. 1 had the largest number of meiotic *24-PHAS* loci (Figure S3).

For both premeiotic and meiotic *24-PHAS* loci, the major sRNAs were enriched for a 5’-terminal adenine (5’-A). For the *21-PHAS* loci, in all eight varieties, the 19^th^ nucleotide position of the major sRNAs was consistently enriched for cytosine (C) and underrepresented by A, while the 20^th^ position was enriched for uracil (U) and underrepresented by C. On the other hand, the major sRNAs of the *21-PHAS* loci were enriched for a 5’-A in B73, HP301, NC358, and Oh43, but were enriched for a 5’-C in Il14H, TIL11, TIL25 and RIL003 (Figure S5). Prior work in rice demonstrated that the rice AGO5c [aka MEIOSIS ARRESTED AT LEPTOTENE1 (MEL1)] preferentially loads sRNAs with a 5’-C (5). Whether the differential enrichment of the 5’ end nucleotide of 21-nt phasiRNAs in the *Zea* varieties affects their sorting onto AGO proteins is yet to be determined.

In all *Zea* varieties, the majority of *PHAS* loci overlapped with transposons (Figure 2D; Dataset S4). The majority of transposons that overlapped with *21-PHAS* loci were terminal inverted repeats (TIRs), which are DNA transposons, whereas the majority of those that overlapped with *24-PHAS* loci – both premeiotic and meiotic loci – were long terminal repeats (LTRs), which are retrotransposons (Figures S6A, B, and C). Furthermore, *21-PHAS* overlapped primarily with the PIF/Harbinger-type TIRs, whereas *24-PHAS* overlapped primarily with the Gypsy-type LTRs (Figures S6D, E, and F). These results indicate distinct genomic origins of the 21- and 24-nt phasiRNAs, and suggest that the two size classes of reproductive phasiRNAs may play different roles in transposon silencing (if that is their biological function). Notably, in RIL003, 37% of the *21-PHAS* loci and 32.3% of the meiotic *24-PHAS* loci overlapped with exons (Figure 2D; Dataset S4). These proportions are much higher than in the other varieties, and is possibly due to the fact that the RIL003 transcriptome was assembled and annotated using our anther data and with different methods than those used for annotating the other *Zea* genomes.

Interrogation of a previously published sRNA-seq data set of a null *dcl5* mutant (11) demonstrated that the majority of both the premeiotic and meiotic *24-PHAS* loci were downregulated substantially in *dcl5* (Figure 3), suggesting that both types of 24-nt phasiRNAs are dependent on DCL5. In support of this notion, *DCL5* is expressed in the premeiotic anthers of wild-type maize, although at lower levels compared to the 1.5-mm stage (Figure S7) (22). Notably, many *24-PHAS* loci of both types still produce detectable levels of 24-nt phasiRNAs in *dcl5* (Figures 3A and 3B), indicating that *24-PHAS* precursors are processed by another DCL protein in the absence of DCL5. This DCL protein is possibly DCL3, a paralog of DCL5 and known to process 24-nt siRNAs (2). Moreover, the downregulation of meiotic 24-nt phasiRNAs was significantly more substantial than the premeiotic 24-nt phasiRNAs at both 1.5- and 2-mm stages (*P* < 0.05; Figure 3C), suggesting that the meiotic 24-nt phasiRNAs are more crucially dependent on DCL5 than the premeiotic 24-nt phasiRNAs. Using RNA-seq data from the prior DCL5 study (11), we performed a differential transposon expression analysis of 1.5-mm anthers from the *dcl5* mutant, but did not detect any differentially expressed transposons [fold change (FC) > 1.5, FDR < 0.05; Dataset S5], suggesting that both classes of 24-nt phasiRNAs are unlikely to transcriptional repress transposons.

**Figure 3.**
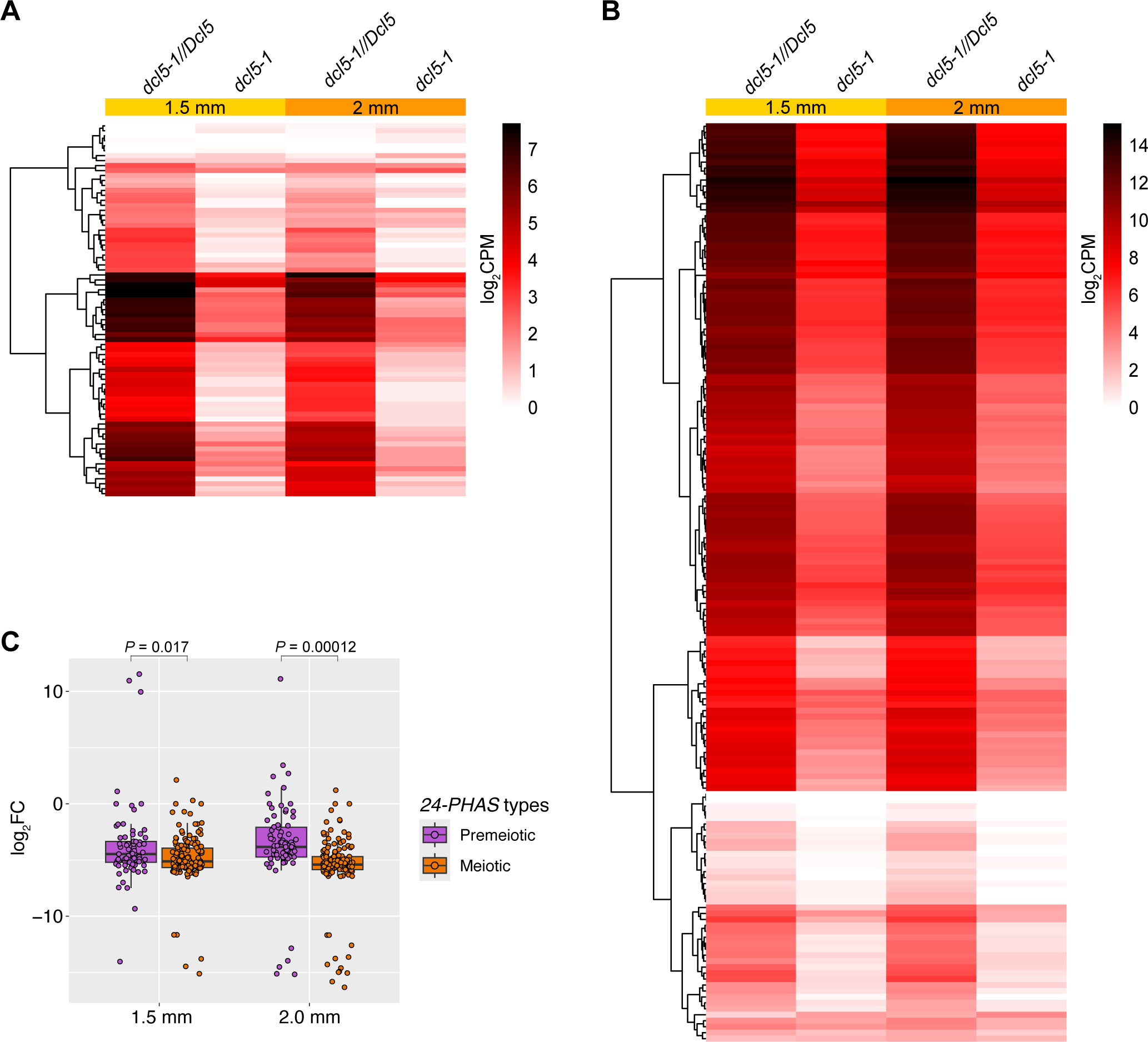
Abundance of premeiotic and meiotic 24-nt phasiRNAs in the previously reported *dcl5-1* mutant compared to heterozygous siblings (*dcl5-1//Dcl5*) (11). (**A and B**) Abundance of 24-nt sRNAs derived from premeiotic (A) and meiotic (B) *24-PHAS* loci. In both heatmaps, each row represents a *24-PHAS* locus. Heatmaps were clustered on Euclidean distance. (C) Box plot of log2FC of the abundance of 24-nt phasiRNAs derived from premeiotic and meiotic *24-PHAS* loci. The *P* values were calculated using unpaired Student’s *t* test.

Premeiotic 24-nt phasiRNAs have been reported in barley and wheat, but not in maize or rice (18). Thus, we identified *24-PHAS* loci in rice and examined their temporal expression patterns using two previously published sRNA-seq data sets (3, 23) using the same methods we applied to our *Zea* data. We found that 34 of the 79 *24-PHAS* loci in rice already produced abundant phasiRNAs at premeiotic stages (Figure S8; Dataset S6). Similar to the *Zea* varieties, the rice premeiotic *24-PHAS* loci are significantly longer than rice *21-PHAS* and meiotic *24-PHAS* loci (Figure S9). Therefore, we conclude that premeiotic 24-nt phasiRNAs are more broadly present in grasses than previously thought.

### Identification of miRNA loci in the *Zea* genomes

Using ShortStack (24), miR-PREFeR (25), and miRador (26), we annotated the expressed miRNAs in the *Zea* genomes. A union of miRNAs identified by the three methods yielded 175 (in NC358) to 366 (in TIL11) mature miRNA sequences encoded by each genome (Figure S10 and Dataset S7). In every variety, we detected expression of at least one copy of *miR2118* and *miR2275* in the anthers (Dataset S8), consistent with their canonical roles in triggering reproductive phasiRNAs. To identify all putative *MIR2118* and *MIR2275* loci in the *Zea* genomes, including those without detectable mature miRNAs in our anther samples, we employed a BLAST-based approach to identify all putative *MIR2118*/*MIR2275* loci in the eight genomes, without relying on sRNA-seq reads. This led to identification of 15 to 17 *MIR2118* loci and 6 to 11 *MIR2275* loci in each genome (Dataset S9). The numbers of *MIR2118/MIR2275* copies and their chromosomal distributions are similar across the *Zea* varieties suggesting high levels of functional conservation of the two miRNA families in the *Zea* genus.

### The premeiotic 24-nt phasiRNAs are unique in triggering mechanism and targets

To identify the triggers of reproductive phasiRNAs in the *Zea* varieties, we predicted interactions between miR2118 and *21-PHAS* precursors and between miR2275 and *24-PHAS* precursors using sPARTA (27). In each *Zea* variety, 64.1% to 84.7% of the *21-PHAS* precursors have a predicted miR2118 target site, and 66.1% to 79.0% of the meiotic *24- PHAS* have a predicted miR2275 target site, respectively. In contrast, only 3.4% to 17.0% of the premeiotic *24-PHAS* have a predicted miR2275 target site (Dataset S10). Consistent with these predicted miRNA-*PHAS* interactions, an enrichment analysis of ∼22-nt motifs in the *PHAS* loci of each *Zea* variety detected a miR2118-matching motif in the *21-PHAS* loci and a miR2275-matching motif enriched in the meiotic *24-PHAS* loci, whereas no miR2275-like motif was found to be enriched in the premeiotic *24-PHAS* loci (Dataset S11). Moreover, none of the 20- to 22-nt motifs enriched in the premeiotic *24-PHAS* loci were similar to known maize miRNAs in sequences (Dataset S11), suggesting that the maize premeiotic 24-nt phasiRNAs are not triggered by miRNAs.

To validate the phasiRNA triggers in the *Zea* varieties, we performed nanoPARE (28) analyses on the same anther materials we used for sRNA-seq. In line with our predictions above, many *21-PHAS* precursors and meiotic *24-PHAS* precursors were targeted by miR2118 or miR2275 (Dataset S12, Sheets 1 and 3–22). In TIL11, we observed the largest number of miR2118–*21-PHAS* interactions at 0.7 mm compared to the other stages, and the largest number of miR2275–*24-PHAS* at 1.5mm (Dataset S12, Sheet 1), indicating stage-specific biogenesis of reproductive phasiRNAs. Notably, premeiotic *24-PHAS* precursors were rarely targeted by miR2275 or any other known miRNAs based on our nanoPARE analysis (Dataset S12, Sheet 1). This result confirms that the premeiotic 24-nt phasiRNAs are likely not triggered by miRNAs.

Using the nanoPARE data, we also detected several miRNA-mediated mRNA cleavage events in each *Zea* variety (Dataset S12, Sheet 2). Notably, several of the miRNA families mediate cleavage of mRNAs encoded by orthologous genes in several of the *Zea* genomes. For example, miR160, miR171, and miR396 each regulates one or two genes that are highly conserved (i.e., within a pangene set) across all *Zea* varieties with available pangene annotation (i.e., except RIL003), plus several other genes that are in a pangene set comprising fewer *Zea* genomes. The highly conserved miRNA-target pairs are likely to play a key role in regulating anther development.

An analysis of phasiRNA targets showed that, in every *Zea* variety, phasiRNAs regulate a small number of protein-coding transcripts, and only a few targets of the meiotic 24-nt phasiRNAs belonging to two pangene sets are conserved in three or four *Zea* varieties (Dataset S13, Sheet 2). These results suggest that either the majority of reproductive phasiRNAs function via a mechanism other than mediating mRNA cleavage or they act on only a small number of target genes that diverged rapidly in the *Zea* genus. Consistent with a prior study in maize and rice (12), 21-nt phasiRNAs can mediate *cis*-cleavage in all *Zea* varieties (Dataset S13, Sheets 1 and 3–22). Interestingly, our analyses also detected many *21-PHAS* precursors that are targeted by 21-nt phasiRNAs *in trans* (Dataset S13, Sheet 1).

Furthermore, meiotic 24-nt phasiRNAs also mediate *cis*-cleavage and/or *trans*-cleavage of other *24-PHAS* precursors in most of the varieties (Dataset S13, Sheet 1). In contrast, the premeiotic 24-nt phasiRNAs do not mediate *cis*- or *trans*-cleavage of *PHAS* precursors in nearly all the varieties with the exception of two *cis*-cleavage events detected in B73). These results suggest that the premeiotic 24-nt phasiRNAs are distinct from the meiotic 24-nt phasiRNAs (and the 21-nt phasiRNAs) in their abilities to mediate *PHAS* precursor cleavage.

### *AGO18* genes are essential for normal accumulation of meiotic 24-nt phasiRNAs but not the premeiotic 24-nt phasiRNAs

AGO18b is known to load 21- and 24-nt phasiRNAs in maize (7), and in rice, the single-copy *AGO18* gene has been shown to be crucial for male fertility (29). To understand the role of the *AGO18* genes in maize anther development and phasiRNA pathways, we generated a triple knockout mutant of all three *AGO18* copies (Figure S1E) by crossing an *ago18a;b* double null mutant that we generated previously using CRISPR-Cas9 (30) with a *ago18c* null mutant obtained from the Mu-Illumina population (31) (Figure S11). The triple homozygous mutant exhibited no obvious defect in fertility, forming fertile tassels and anthers, viable pollen grains, and fully pollinated ears when self-pollinated (Figure S12), suggesting that the *AGO18* genes are dispensable for male and female fertility under normal growth conditions.

Our RNA-seq analyses of developing anthers detected only 47 genes differentially expressed between the triple homozygous mutant plants and their triple heterozygous siblings at 0.5 mm (premeiotic), one gene differentially expressed at 2 mm (meiotic), and none at 5 mm (postmeiotic) (FC > 1.5, FDR < 0.05; Dataset S14), suggesting that loss of *AGO18* genes has a minor impact on the anther mRNA transcriptome. Using sRNA-seq, we detected, at the 0.5- and 2-mm stages, only one *21-PHAS* loci that was differentially expressed between the *ago18* triple mutant and its triple heterozygous siblings, whereas at the 5-mm stage, three *21-PHAS*, two premeiotic *24-PHAS*, and 123 meiotic *24-PHAS* loci were significantly downregulated in the mutant (Figure 4A; Dataset S15). Accordingly, by the 5-mm stage, the total abundance of meiotic 24-nt phasiRNAs were also downregulated dramatically, with average CPM decreasing from 46664.8 to 3987.3 (FC = 11.7; *P* = 3.8 × 10^-3^) (Figure 4B). In contrast, the total abundance of the 21-nt phasiRNAs or premeiotic 24-nt phasiRNAs was not down- or upregulated significantly at any of the three stages (Figure S13). In addition, we detected only three differentially accumulated miRNAs, including a putative miR159 and a putative miR2275 (Dataset S16). Together, these sRNA-seq results suggest that the *AGO18* genes play a key role in the normal accumulation of meiotic 24-nt phasiRNAs and a minor role in that of the 21-nt phasiRNAs, premeiotic 24-nt phasiRNAs, and miRNAs.

**Figure 4.**
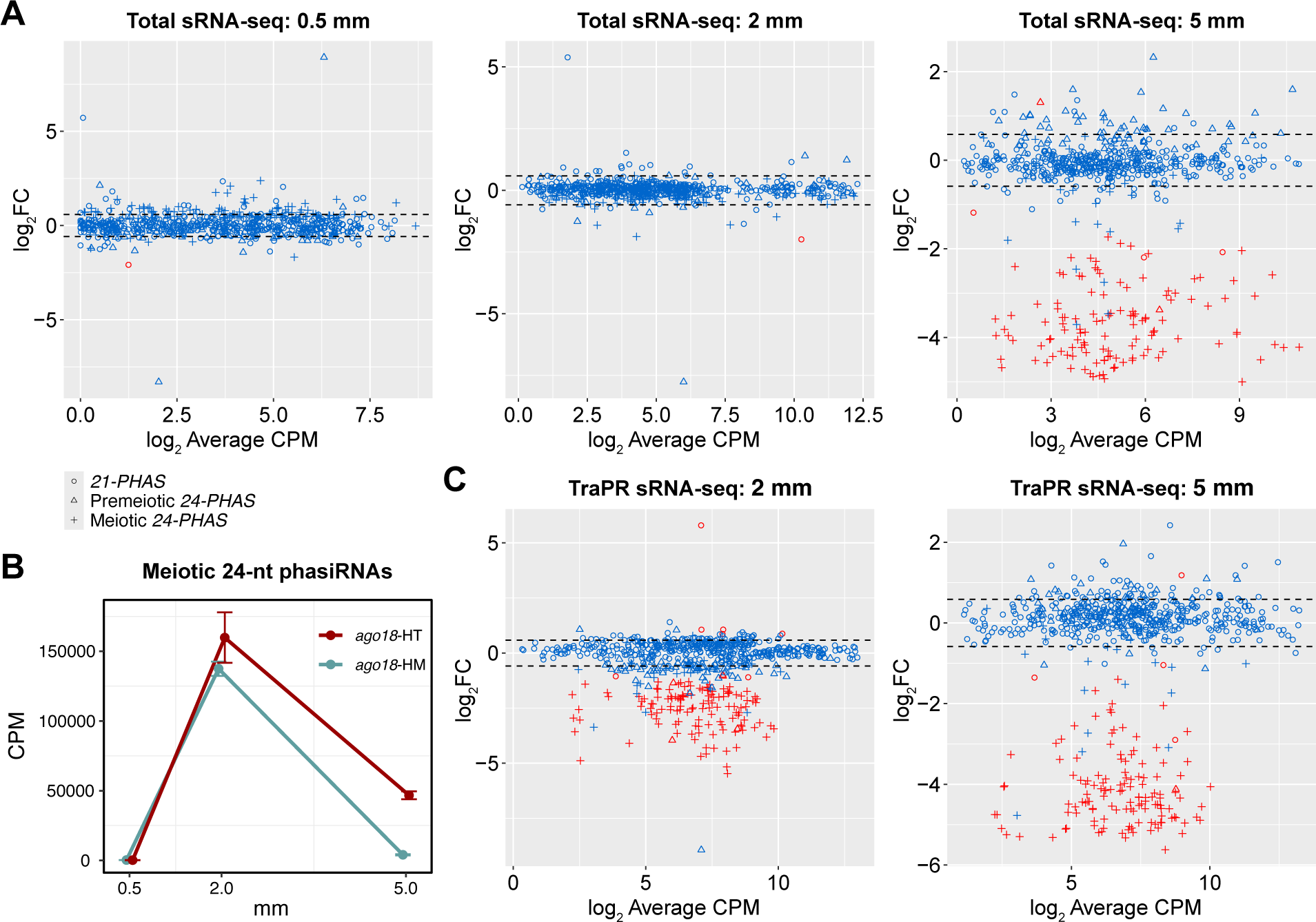
Total sRNA-seq and TraPR sRNA-seq analyses of the *ago18* triple mutant anthers. **(A)** Mean-difference (MA) plots of *PHAS*loci based on sRNA-seq. **(B)** Total abundance (mean ± se) of meiotic 24-nt phasiRNAs in *ago18* triple homozygous mutant plants (*ago18*-HM) and their triple heterozygous siblings (*ago18*-HT). Results of Student’s *t* test are shown in Figure S13. **(C)** MA plots from differential accumulation analyses of phasiRNA abundance per loci based on TraPR sRNA-seq. In (A) and (C), average CPM values were calculated for only triple heterozygous samples. Significantly differentially expressed loci (FC > 1.5, FDR &lt; 0.05) are represented by red circles/triangles/plus signs, and the other loci in blue. Dash lines indicate FC = ±1.5 (i.e., log2FC = ±0.585).

To characterize the impact of loss of *AGO18* genes on the profile of AGO-loaded sRNAs, we carried out high-throughput sequencing of sRNAs isolated using the TraPR method (32) from 2- and 5-mm anthers of the *ago18* triple mutant. Interestingly, in contrast to the total sRNA-seq results described above, 98 meiotic *24-PHAS* loci, two premeiotic *24- PHAS* loci, and two *21-PHAS* loci were already downregulated in phasiRNA abundance at 2 mm. At the 5-mm stage, 129 meiotic *24-PHAS* loci, one premeiotic *24-PHAS*, and four *21- PHAS* loci were downregulated in the mutant (Figure 4C; Dataset S17). Moreover, the total abundance of meiotic 24-nt phasiRNAs demonstrated significant downregulation at 2 and 5 mm, whereas the premeiotic 24-nt phasiRNAs did not show this (Figure S14). Together, our total sRNA-seq and TraPR sRNA-seq analyses indicate that, at 2 mm, the loss of *AGO18* genes does not affect the abundance of meiotic 24-nt phasiRNAs, but significantly fewer meiotic 24-nt phasiRNAs are loaded onto AGOs, suggesting that AGO18 proteins are essential for loading meiotic 24-nt phasiRNAs but not for their accumulation at the 2-mm stage. In contrast, at the 5-mm stage, AGO18 proteins are essential for both loading and accumulation of 24-nt phasiRNAs.

NanoPARE analyses of 0.5-, 2-, and 5-mm anthers of the *ago18* triple mutant demonstrated that for both miRNAs and reproductive phasiRNAs, the sRNA-target interactions were consistently fewer in the mutant compared to their heterozygous siblings (Dataset S18), suggesting that AGO18 facilitates miRNA and phasiRNA-mediated gene silencing. Furthermore, at 0.5 and 2 mm, we detected many *trans*-cleavage events of *PHAS* precursors by 21-nt and meiotic 24-nt phasiRNAs, consistent with our analysis of wild-type maize varieties, yet most of the cleavage events were not detected in the triple mutant (Dataset S18), suggesting that the loading of 21-nt phasiRNAs and meiotic 24-nt phasiRNAs are crucial for *cis*- and *trans*-cleavage of reproductive *PHAS* precursors.

## DISCUSSION

We demonstrate that the premeiotic 24-nt phasiRNAs are derived from genomic loci that are distinct from the *21-PHAS* or meiotic *24-PHAS* loci and are present in all the *Zea* varieties that we analyzed – including five maize inbred lines and three teosinte species/subspecies (Figure 1) – and rice. In the *Zea* genus, premeiotic 24-nt phasiRNAs exhibited several similarities and differences compared to meiotic 24-nt phasiRNAs. The two types of 24-nt phasiRNAs are similar in that (i) their genomic origins, which overlap predominantly with transposons (Figure 2D and Figure S6); and (ii) their dependence on DCL5 for biogenesis (Figure 3). However, the premeiotic 24-nt phasiRNAs are unique in several aspects. First, their genomic loci are distinct and significantly longer than those of the 21-nt and meiotic 24- nt phasiRNAs (Figure 2C). Second, they are likely not triggered by miRNA-mediated cleavage of *PHAS* precursors, but rather by an unknown mechanism (Dataset S11). Third, the vast majority of premeiotic 24-nt phasiRNAs are likely not loaded by AGO18, whereas many meiotic 24-nt phasiRNAs are (Figure 4). Fourth, while both the 21-nt phasiRNAs and the meiotic 24-nt phasiRNAs can mediate *cis*-cleavage, the premeiotic 24-nt phasiRNAs generally do not mediate *cis*-cleavage of their precursors (Tables S13). Together, these findings indicate that the premeiotic 24-nt phasiRNAs constitute a unique class of reproductive phasiRNAs with respect to both biogenesis mechanism and molecular function. We speculate that a possible reason that the premeiotic 24-nt phasiRNAs were not identified in previous work (1) was that the anther sRNA data were generated from the W23 *bz2* inbred line, but were aligned to the B73 reference genome for analyses, requiring zero mismatch in identifying *PHAS* loci. Thus, mapping sRNA data to the corresponding genome might be crucial; this is easier today than it was in 2015 because of the proliferation of high quality genomes.

Our observations that only a small number of anther-expressed protein-coding genes are downregulated in the *ago18* triple mutant (Dataset S14) but many *24-PHAS* loci are downregulated in sRNA abundance (Figures 4A; Dataset S15) suggest that in the anthers the *AGO18* genes function predominantly in the reproductive phasiRNA pathways. The substantial differences we observed in total sRNA-seq versus TraPR sRNA-seq analyses of the *ago18* mutant – especially at the 2-mm stage (Figures 4A and 4C) – indicate that the meiotic 24-nt phasiRNAs can accumulate in maize anthers without being loaded onto an AGO protein, at least for a period of time. However, by the post-meiotic phase of anther development, the AGO18 proteins seem necessary for their normal accumulation (Figure 4A). This in turn suggests that the meiotic 24-nt phasiRNAs may have crucial functions during post-meiotic anther development. Nonetheless, the lack of an obvious phenotype of the *ago18* triple mutant (Figure S12), contradicting a prior report showing that *ago18* is crucial for male fertility (29), suggests that perhaps in maize other AGO protein(s) act redundantly or in a compensatory manner with AGO18 in the phasiRNA pathways.

In summary, this work demonstrates that the premeiotic 24-nt phasiRNAs are more broadly present – at least in the grass lineage – than previously thought. Our data provide significant insights into the biogenesis and molecular function of the premeiotic 24-nt phasiRNAs, including the roles of several biogenesis factors or effector proteins such as DCL5 and AGO18. However, the identity of the other key players in the premeiotic 24-nt phasiRNA pathway, and whether premeiotic 24-nt phasiRNAs exist in other grass species and more broadly across monocots, remains unknown. Therefore, more work will be needed to further understand the biogenesis, function, and evolution of these novel premeiotic 24-nt phasiRNAs.

## MATERIALS AND METHODS

### Plant materials and growth

Seeds of HP301, Il14H, NC358, Oh43 and TIL11 were provided by Matthew Hufford (Iowa State University), TIL25 seeds were provided by John Doebley (University of Wisconsin-Madison), and RIL003 seeds were obtained through the U.S. National Plant Germplasm System (accession: PI 422162). The *ago18* triple mutant was generated by crossing a previously reported *ago18a;b* double mutant (30) with an *ago18c* mutant obtained from the Mu-Illumina population (insertion ID: mu-illumina_50570.7) (31) ; the *ago18* materials analyzed in this work are transgene free. Maize plants were grown in greenhouses of the Donald Danforth Plant Science Center Plant Growth Facility under 14-hour day length, 28°C/24°C temperature cycles, and 50% humidity, and the teosinte varieties were grown in a walk-in growth chamber at the same facility under 12-hour day length, 28°C/23°C temperature cycles, and 40% humidity.

### Cytological analysis

Fresh anthers were fixed with 2% [v/v] paraformaldehyde, 2% [v/v] glutaraldehyde, and 0.1% [v/v] Tween-20 in 0.1 M PIPES buffer (pH 7.4) overnight, dehydrated using a concentration gradient of acetone (30%, 50%, 70%, 80%, 90%, and 100% [v/v]), embedded using a Quetol 651 - NSA Kit (no. 14640, Electron Microscopy Sciences), and polymerized at 60°C. Embedded tissues were sectioned into 500 nm sections using a Leica Ultracut UCT (Leica Microsystems Inc., Wetzlar, Germany), and stained using the Epoxy Tissue Stain solution (no. 14950, Electron Microscopy Sciences). Anther sections were imaged using a Leica DM 750 microscope. Images were captured with a Leica ICC50 HD camera and Leica Acquire v2.0 software (Leica Microsystems Inc., Wetzlar, Germany).

### Library preparation and sequencing

Fresh anther samples, each with 2 or 3 biological replicates derived from distinct plants, were snap-frozen in liquid nitrogen and total RNA was extracted using the TRI reagent (Sigma-Aldrich) or the TraPR Small RNA Isolation kit (Lexogen; for TraPR sRNA-seq only). sRNA-seq libraries were prepared from ∼100 ng total RNA per sample using a Somagenics RealSeq-AC miRNA library kit following the manufacturer’s protocol. RNA-seq (Smart-seq2) and nanoPARE libraries were prepared from 5 ng total RNA per sample using the nanoPARE library preparation protocol (28) with previously described modifications (17). All libraries were sequenced on an Illumina NextSeq 550 instrument at the University of Delaware DNA Sequencing & Genotyping Center to generate 76-nt single-end reads.

### Whole proteome-based phylogenetic analysis

The anther transcriptome of RIL003 was assembled from the RNA-seq and nanoPARE reads. Briefly, StringTie v2.1.7 (33) and Scallop v0.10.4 (34) were separately used with default parameters to perform *de novo* transcriptome assembly, and the resulting transcriptome annotations were merged using the merge function of StringTie. Protein-coding transcripts were identified using TranSuite v0.2.2 (35) with parameter “Auto”. RIL003 transposons were annotated using Extensive de-novo TE Annotator (EDTA) v2.0.1 (36) with parameters --species Maize --anno 1 --force 1. Annotation of gene models and transposons of all the other *Zea* genomes (B73 RefGen_v5 and v1 of all the other varieties) were obtained from maizeGDB (37). Whole proteome sequences of the outgroup species were obtained from Ensembl Plants (*Chondrus crispus*) or Phytozome v13 (all the other species). A species tree was generated using Orthofinder v2.5.4 (38) with default parameters. Gene-family trees were built using previously described methods (39).

### *PHAS* loci identification and analyses

*PHAS* loci were identified from sRNA-seq data using ShortStack v3.8.5 with parameters -- mismatches 0 --mincov 0.5rpm and phasing scores ≥ 30 as the cutoff. The length of each *PHAS* precursor, as estimated by ShortStack, was used as a proxy to the length of the *PHAS* locus. Premeiotic *24-PHAS* loci were defined as those with a minimal CPM (mean of replicates) > 20 at premeiotic stages, and the remaining *24-PHAS* loci were defined as meiotic. Rice sRNA-seq were obtained from two previous studies (3, 23). Data from (3) were used to identify *PHAS* loci, and data from both studies were separately normalized and analyzed for abundance of 24-nt phasiRNAs per loci. Premeiotic *24-PHAS* loci were defined as those with minimal CPM > 20 at premeiotic stages based on data from (23). Overlap among the three types of *PHAS* loci or between *PHAS* loci and various genomic features were determined using the intersect function of BEDtools v2.29.2 (40) with parameters -e -f 0.5 -F 0.5.

### microRNA loci identification and analyses

To annotate expressed miRNA loci, all sRNA-seq data for each genotype were analyzed using ShortStack v3.8.5 with parameters --mismatches 0 --dicermax 22 --mincov 15, miR- PREFeR v0.24 with parameters PRECURSOR_LEN = 300, READS_DEPTH_CUTOFF = 20, MIN_MATURE_LEN = 20, MAX_MATURE_LEN = 22, ALLOW_NO_STAR_EXPRESSION = N, ALLOW_3NT_OVERHANG = N, CHECKPOINT_SIZE = 300, and miRador with default parameters. Using BLASTN, miRNAs were assigned to known miRNA families in miRBase (41) release 22.1 (≤ 5 mismatches) or defined as novel miRNAs (requiring corresponding miRNA* reads). For each genome, a union of all the expressed mature miRNAs identified using the three tools were used for downstream analyses.

To identify all putative *MIR2118* and *MIR2275* loci in the *Zea* genomes, publicly available *MIR2118*/*MIR2275* precursor sequences were obtained from miRBase and a previous study (42) and used as queries to search for homologous sequences in the *Zea* genomes using BLASTN with parameters -max_target_seqs 10 -evalue 10 -word_size 10. The subject regions that were longer than 80% of the length of query sequences were filtered and merged using the merge function of BEDtools. The resulting sequences were aligned using MUSCLE (43), and phylogenetic trees were built using IQ-TREE v2.2.0.3 (44) to assign/curate names of *MIR2118* and *MIR2275* loci based on orthology. MUSCLE-generated alignments were examined using Jalview v2 (45) to identify putative mature miR2118/miR2275 sequences and corresponding miRNA* sequences that are homologous to known members of the families.

### sRNA/mRNA quantification and differential expression analyses

To quantify phasiRNA abundance, sRNA-seq reads were mapped to the respective *Zea* genomes using Bowtie v1.3.1 (46); to quantify miRNA abundance, sRNA-seq reads were mapped to a FASTA file of all mature miRNAs annotated for each genome using Bowtie; and to quantify mRNA abundance, RNA-seq/Smart-seq2 reads were mapped to the respective *Zea* genomes (B73 RefGen_v5 was used for the *dcl5* and *ago18* mutant data) using HISAT v2.1.0 (47) with parameters --min-intronlen 30 --max-intronlen 8000 . Reads mapped to *PHAS* loci or miRNAs were counted using featureCounts v1.6.3 (48) with parameter -M and normalized to counts per million (CPM) using edgeR v4.0.2 (49). Differential expression analyses were performed using edgeR with a generalized linear model-based method.

### Small RNA target identification

Small RNA-transcript interactions in each anther sample were identified using the nanoPARE analysis pipeline (28). The sRNA-target pairs with adjusted *P* value < 0.05 at the EndCut step and are detected in at least two biological replicates were considered positive.

### Identification of RNA polymerase II-derived strands of *PHAS* loci

NanoPARE reads mapped to both strands of genomic DNA were counted separately using featureCounts for each *PHAS* locus. For loci with >10 raw reads, the RNA strand that accounts for >70% of the total reads were considered as the RNA polymerase II-derived strand of a precursor.

### Motif enrichment analysis

For detection of potential miRNA target sites in *PHAS* precursors, *PHAS* loci were extended by 500 bp on both ends. The sequences of extended *PHAS* loci were obtained using the getfasta function of BEDTools, and motif enrichments were detected using the MEME program (*E* value < 0.05) of the MEME Suite (50). Enriched motifs were aligned with known maize miRNA sequences from miRBase using the Tomtom program (*E* value < 0.001) of the MEME Suite.

### Quantification of male fertility

Anther exertion was monitored daily, and tassels were detached on the day when anthers in the lower florets of the lowest tassel branch exerted. RGB images of the tassels were captured using a digital single-lens reflex camera controlled by a Raspberry Pi within an enclosed light tent to ensure consistent imaging settings. Four photos were taken for each tassel with 90-degree rotations to capture variations from different angles. The Tasselyzer pipeline (51) was applied to the tassel images to quantify anther exertion. Images were analyzed at the pixel level based on color information and segmented into anthers, other tassel areas, and background, using a naïve Bayes classifier on the PlantCV platform (52). The ratio of anther pixels to the total pixels of anther and other tassel areas was calculated to quantify male fertility mimicking visual observations. Anther/tassel area ratios were averaged among the four images taken from different angles. Fuchsia and green highlights were applied to anthers and other tassel areas for visual inspection of segmentation results and demonstration purposes.

### Pollen imaging

Anthers were dissected shortly before anthesis and placed in a 1.7 mL tube. About 6 anthers were stained in 35 µl of Alexander staining solution for 5 minutes. During staining, anthers were gently squeezed using forceps to release pollen grains. The tube was centrifuged at 100x *g* for 1 min, and the debris of somatic cells were removed using forceps. Pollen grains were washed twice in 100 µL of 1x PBS buffer, resuspended in 35 µL of 50% glycerol solution, mounted on a microscope slide, and imaged using a Leica DM 750 microscope as described above for the cytological analysis of anthers.

## ACKNOWLEDGMENTS

This work was supported by the National Key R&D Program of China (award 2023ZD04073, to J.Z.), the National Institute of General Medical Sciences of the US National Institutes of Health (award 1R01GM151302-01, to B.C.M.), and the US National Science Foundation (award 1754097, to B.C.M.). We thank Matt Hufford for providing seeds of the maize inbred lines HP301, Il14H, NC358, and Oh43 and teosinte TIL11, John Doebley for providing teosinte TIL25, Brewster Kingham and Mark Shaw (University of Delaware DNA Sequencing & Genotyping Center) for assistance with Illumina sequencing, Mayumi Nakano for assistance with data handling, and Joanna Friesner for assistance with editing.

## AUTHOR CONTRIBUTIONS

J.Z. and B.C.M. designed research; J.Z., S.B., S.L., C.T., and M.M. performed research; A.B., M.A.S., and M.D.N. contributed new analytic tools; J.Z. and B.C.M. analyzed data and wrote the paper.

## SUPPLEMENTAL DATA

Figure S1. Maximum-likelihood phylogeny of small RNA pathway-associated protein families HD-ZIP IV (A), bHLH (B), RDR (C), DCL (D), AGO (E), SGS3 (F), DRB (G), SE (H), SDN (I), and HESO1/URT1 (J) in the *Zea* varieties and outgroup species shown in Figure 1A. The HD-ZIP IV clade was pruned from a bigger homeobox (HB) family phylogeny, and in the

bHLH phylogeny, clades that are distantly related to the MS23/MS32/bHLH51/bHLH122 were collapsed.

Figure S2. Temporal accumulation patterns of phasiRNAs derived from individual *21-PHAS* loci in each *Zea* variety. Heatmaps are color-coded by log2-transformed average CPM of biological replicates.

Figure S3. Numbers of *PHAS* loci on individual chromosomes of each *Zea* variety.

Figure S4. Chromosomal distribution of *PHAS* loci in each *Zea* variety. *21-PHAS*, premeiotic *24-PHAS*, and meiotic *24-PHAS* loci are represented by dots with the same color scheme as Figures 2B and 2C. Height of dots corresponds to log2-transformed maximal CPM (Dataset S2).

Figure S5. Nucleotide composition of 21-nt phasiRNAs, premeiotic 24-nt phasiRNAs, and meiotic 24-nt phasiRNAs in each *Zea* variety.

Figure S6. Percentages of *PHAS* loci that overlap with different transposon types in each *Zea* variety. In panels A–C, transposons were separated based on the major categories, whereas in D–F, transposons were classified into more specific subtypes.

Figure S7. Temporal expression pattern of *Dcl5* in wild-type maize (W23 *bz2* inbred line) anthers and pollen. Numbers on the *x*-axis are developmental stages represented by anther lengths (in mm). Normalized RNA-seq data in reads per kilobase of transcript per million reads mapped (RPKM) were obtained from a prior publication (22).

Figure S8. sRNA abundance of rice *24-PHAS* loci based on previously published data. *24-PHAS* loci were identified using the raw sRNA-seq data from (3, 23). Data from the two studies were separately normalized. In rice anther development, it had been demonstrated that stages before stage 7 (S7) are premeiotic, S7–S8 are meiotic, while S9 and later stages are postmeiotic (54).

Figure S9. Boxplot of lengths of *PHAS* loci in rice. *P* values were calculated using one-way ANOVA with post-hoc Tukey’s HSD test.

Figure S10. Venn diagrams of mature miRNAs identified by miRaor, ShortStack, or miR-PREFeR in each *Zea* variety.

Figure S11. Nature of the *ago18a/b/c* mutations in the maize *ago18* triple mutant. The *ago18a* and *ago18b* mutations are single-nucleotide deletion in the coding DNA sequences, causing frameshift and premature stop codons, whereas the *ago18c* mutant allele has a Mu transposon insertion in the coding sequence. CRISPR guide RNA sequences are shown in magenta, and the adjacent protospacer adjacent motif (PAM) sequences in green. Underscore indicates an intron in the *AGO18b* sequence.

Figure S12. Phenotypic analyses of the maize *ago18* triple mutant. (A and B) Quantification of male fertility using the Tasselyzer method. Tassels of 5 triple homozygous (*ago18*-HM) and 6 triple heterozygous siblings (*ago18*-HT) were phenotyped and one representative tassel of each genotype is shown in (A). The *P* value in (B) was calculated using unpaired Student’s *t* test. (C) Micrographs of mature pollen grains treated with Alexander staining solution. Pollen from 7 triple homozygous and 9 triple heterozygous siblings were examined and one representative image of each genotype is shown. (D) Images of self-pollinated ears. Ears from 5 triple homozygous and 8 triple heterozygous siblings were examined and one representative image of each genotype is shown.. Scale bars: (A) and (E), 2 cm; (C), 20 μm; (D), 200 μm.

Figure S13. Total abundance of each phasiRNA class in the *ago18* triple homozygous mutant plants (*ago18*-HM) versus their triple heterozygous siblings (*ago18*-HT) based on total sRNA-seq. **(A)** Total abundance (mean ± se) of 21-nt phasiRNAs (*Left*) and premeiotic 24-nt phasiRNAs (*Right*). **(B)** *P* values of the differences in phasiRNA abundance between *ago18*- HM and *ago18*-HT calculated using unpaired Student’s *t* test. The only *P* value smaller than 0.05 is in bold and red.

Figure S14. Abundance of AGO-loaded sRNAs of each phasiRNA class in the *ago18* triple homozygous mutant plants (*ago18*-HM) versus their triple heterozygous siblings (*ago18*-HT) based on TraPR sRNA-seq. **(A)** Abundance (mean ± se) of premeiotic 24-nt phasiRNAs (*Top left*), meiotic 24-nt phasiRNAs (Top right), and 21-nt phasiRNAs (*Bottom left*). **(B)** *P* values of the differences in phasiRNA abundance between *ago18*-HM and *ago18*-HT calculated using unpaired Student’s *t* test. *P* values smaller than 0.05 are in bold and red.

**Dataset S1.** Copy numbers of genes in specific clades of the phylogenies in Figure S1.

**Dataset S2.** Annotation and phasiRNA abundance of reproductive *PHAS* loci in each *Zea* variety.

**Dataset S3.** *P* values of one-way ANOVA with post-hoc Tukey’s HSD test of the lengths of *21-PHAS*, premeiotic *24-PHAS* loci, and meiotic *24-PHAS* loci in each *Zea* variety.

**Dataset S4.** Overlaps between *PHAS* loci and various genomic features including transposons, exons, introns, and intergenic regions in each *Zea* variety.

**Dataset S5.** Differential expression analysis of transposons using previously published *dcl5* RNA-seq data.

**Dataset S6.** Annotation of rice *24-PHAS* loci based on previously published data.

**Dataset S7.** Numbers of mature miRNAs identified by miRaor, ShortStack, or miR-PREFeR in each *Zea* variety.

**Dataset S8.** Abundance of miRNAs in each *Zea* variety.

**Dataset S9.** Annotation of *MIR2118* and *MIR2275* loci in the *Zea* genomes.

**Dataset S10.** Summary of *PHAS* precursor targeting by miR2118, miR2275, or other miRNAs in each *Zea* variety.

**Dataset S11.** Motif enrichments of *21-PHAS*, premeiotic *24-PHAS*, and meiotic *24-PHAS* loci in each *Zea* variety.

**Dataset S12.** miRNA-target interactions detected by nanoPARE analysis of each *Zea* variety.

**Dataset S13.** phasiRNA-target interactions detected by nanoPARE analysis of each *Zea*

variety.

**Dataset S14.** Differentially expressed genes identified through the RNA-seq analysis of the *ago18* triple homozygous mutant versus triple heterozygous siblings.

**Dataset S15.** Differentially expressed *PHAS* loci identified through the sRNA-seq analysis of the *ago18* triple homozygous mutant versus triple heterozygous siblings.

**Dataset S16.** Differentially expressed miRNA identified through the sRNA-seq analysis of the *ago18* triple homozygous mutant versus triple heterozygous siblings.

**Dataset S17.** Differentially expressed *PHAS* identified through the TraPR sRNA-seq analysis of the *ago18* triple homozygous mutant versus triple heterozygous siblings.

**Dataset S18.** sRNA targets identified by the nanoPARE analysis of the *ago18* triple homozygous mutant and triple heterozygous siblings.

