## Supplemental figures for "Premeiotic 24-nt phasiRNAs are present in the *Zea* genus and unique in biogenesis mechanism and molecular function": Zhan_et_al_supp_figures_2024-03.pdf

A

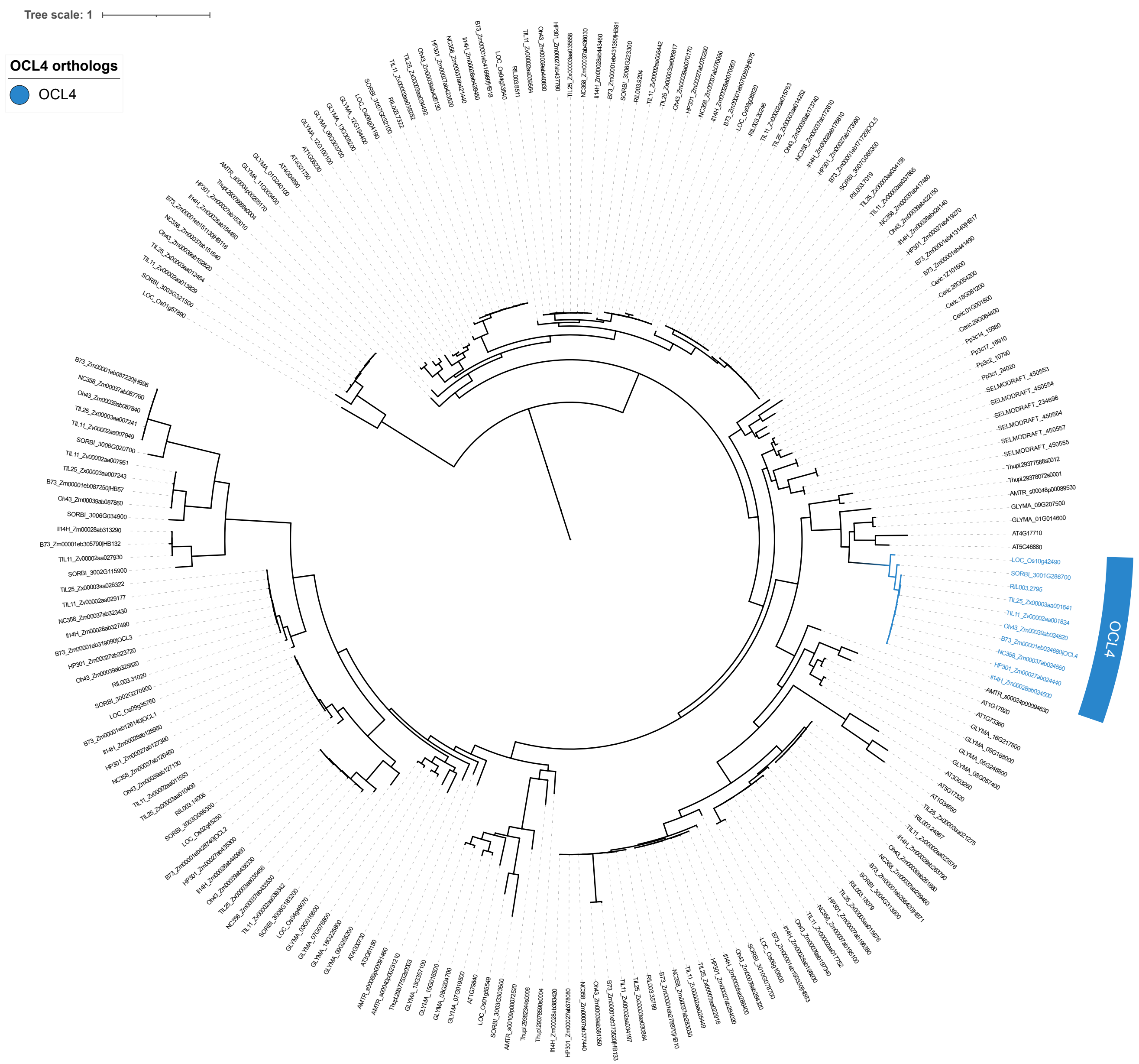


### Figure S1A

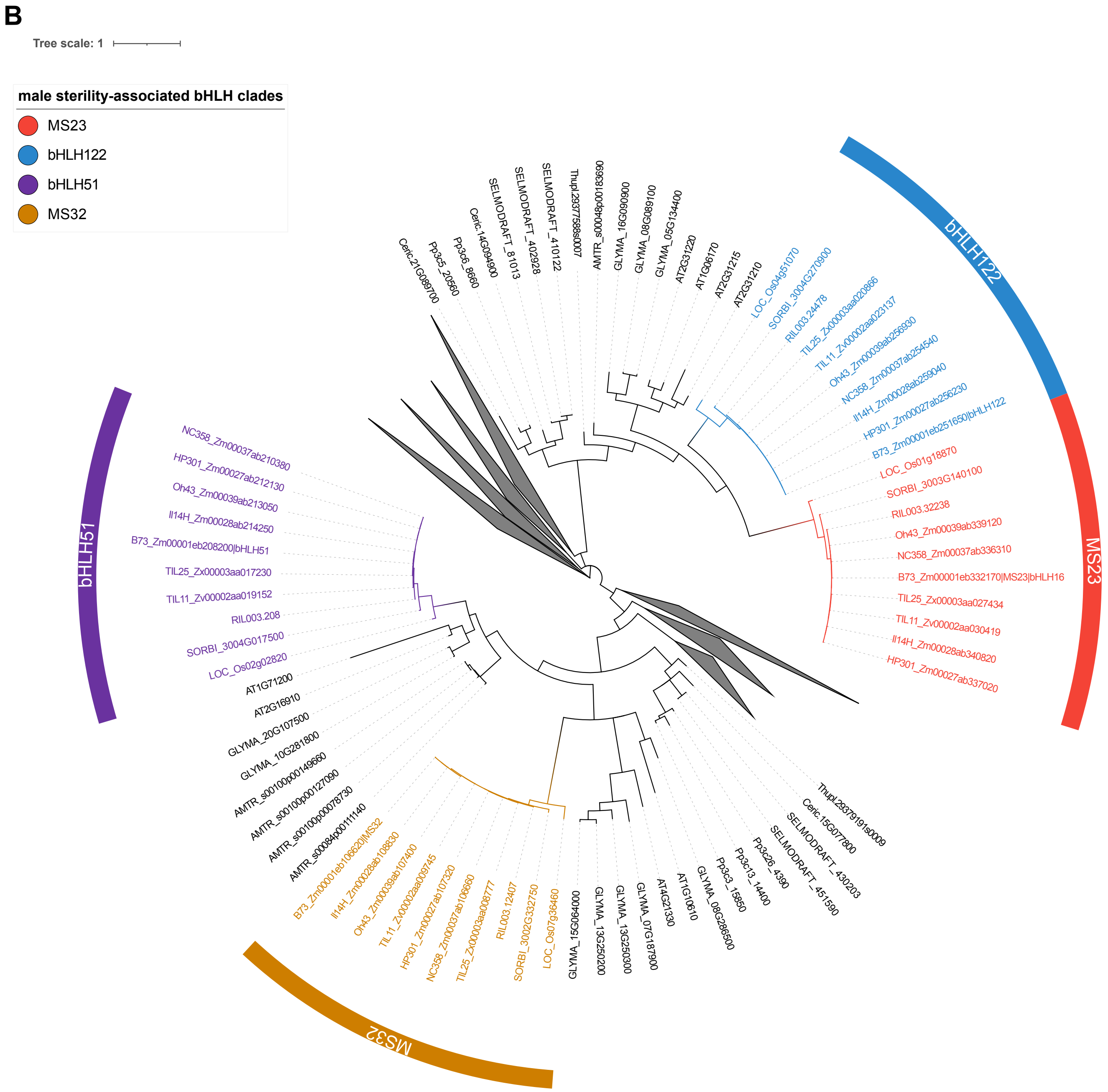
B
Tree scale: 1
male sterility-associated bHLH clades
MS23
bHLH122
bHLH51
MS32

### Figure S1B

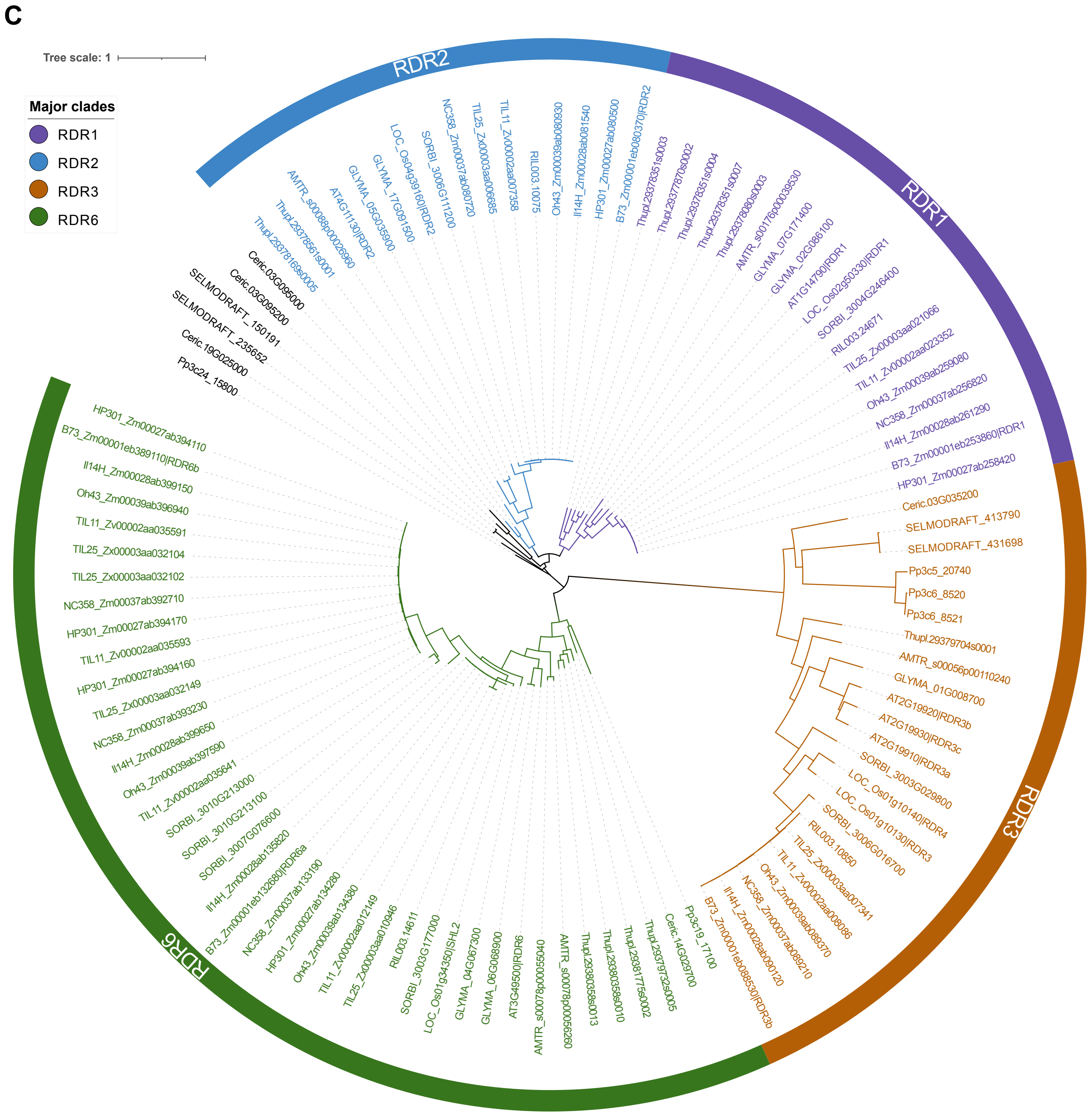

C
Tree scale: 1
Major clades
RDR1
RDR2
RDR3
RDR6
RDR2
RDR1
RDR3
RDR6
Thupl.29378169s0005
Thupl.29378561s0001
AMTR_s00088p00026960
AT4G11130|RDR2
GLYMA_05G035900
GLYMA_17G091500
LOC_Os04g39160|RDR2
SORBI_3006G111200
NC358_Zm00037ab080720
TIL25_Zx00003aa006685
TIL11_Zv00002aa007358
RIL003.10075
Oh43_Zm00039ab080930
Il14H_Zm00028ab081540
HP301_Zm00027ab080500
B73_Zm00001eb080370|RDR2
Thupl.29378351s0003
Thupl.29377870s0002
Thupl.29378351s0004
Thupl.29378351s0007
Thupl.29378080s0003
AMTR_s00176p00039530
GLYMA_07G171400
GLYMA_02G086100
AT1G14790|RDR1
LOC_Os02g50330|RDR1
SORBI_3004G246400
RIL003.24671
TIL25_Zx00003aa021066
TIL11_Zv00002aa023352
Oh43_Zm00039ab259080
NC358_Zm00037ab256820
Il14H_Zm00028ab261290
B73_Zm00001eb253860|RDR1
HP301_Zm00027ab258420
Ceric.03G035200
SELMODRAFT_413790
SELMODRAFT_431698
Pp3c5_20740
Pp3c6_8520
Pp3c6_8521
Thupl.29379704s0001
AMTR_s00056p00110240
GLYMA_01G008700
AT2G19920|RDR3b
AT2G19930|RDR3c
AT2G19910|RDR3a
SORBI_3003G029800
LOC_Os01g10140|RDR4
LOC_Os01g10130|RDR3
SORBI_3006G016700
RIL003.10850
TIL25_Zx00003aa007341
TIL11_Zv00002aa008086
Oh43_Zm00039ab089370
NC358_Zm00037ab089210
Il14H_Zm00028ab090120
B73_Zm00001eb088530|RDR3b
Pp3c19_17100
Ceric.14G029700
Thupl.29379732s0005
Thupl.29381775s0002
Thupl.29380358s0010
Thupl.29380358s0013
AMTR_s00078p00056260
AMTR_s00078p00055040
AT3G49500|RDR6
GLYMA_09G068900
GLYMA_04G067300
LOC_Os01g34350|SHL2
SORBI_3003G177000
RIL003.14611
TIL25_Zx00003aa010946
TIL11_Zv00002aa012149
Oh43_Zm00039ab134380
HP301_Zm00027ab134280
NC358_Zm00037ab133190
B73_Zm00001eb132680|RDR6a
Il14H_Zm00028ab135820
SORBI_3007G076600
SORBI_3010G213100
SORBI_3010G213000
TIL11_Zv00002aa035641
Oh43_Zm00039ab397590
Il14H_Zm00028ab399650
NC358_Zm00037ab393230
TIL25_Zx00003aa032149
HP301_Zm00027ab394160
TIL11_Zv00002aa035593
HP301_Zm00027ab394170
NC358_Zm00037ab392710
TIL25_Zx00003aa032102
TIL25_Zx00003aa032104
TIL11_Zv00002aa035591
Oh43_Zm00039ab396940
Il14H_Zm00028ab399150
B73_Zm00001eb389110|RDR6b
HP301_Zm00027ab394110
Pp3c24_15800
Ceric.19G025000
SELMODRAFT_235652
SELMODRAFT_150191
Ceric.03G095200
Ceric.03G095000


Figure S1C

D

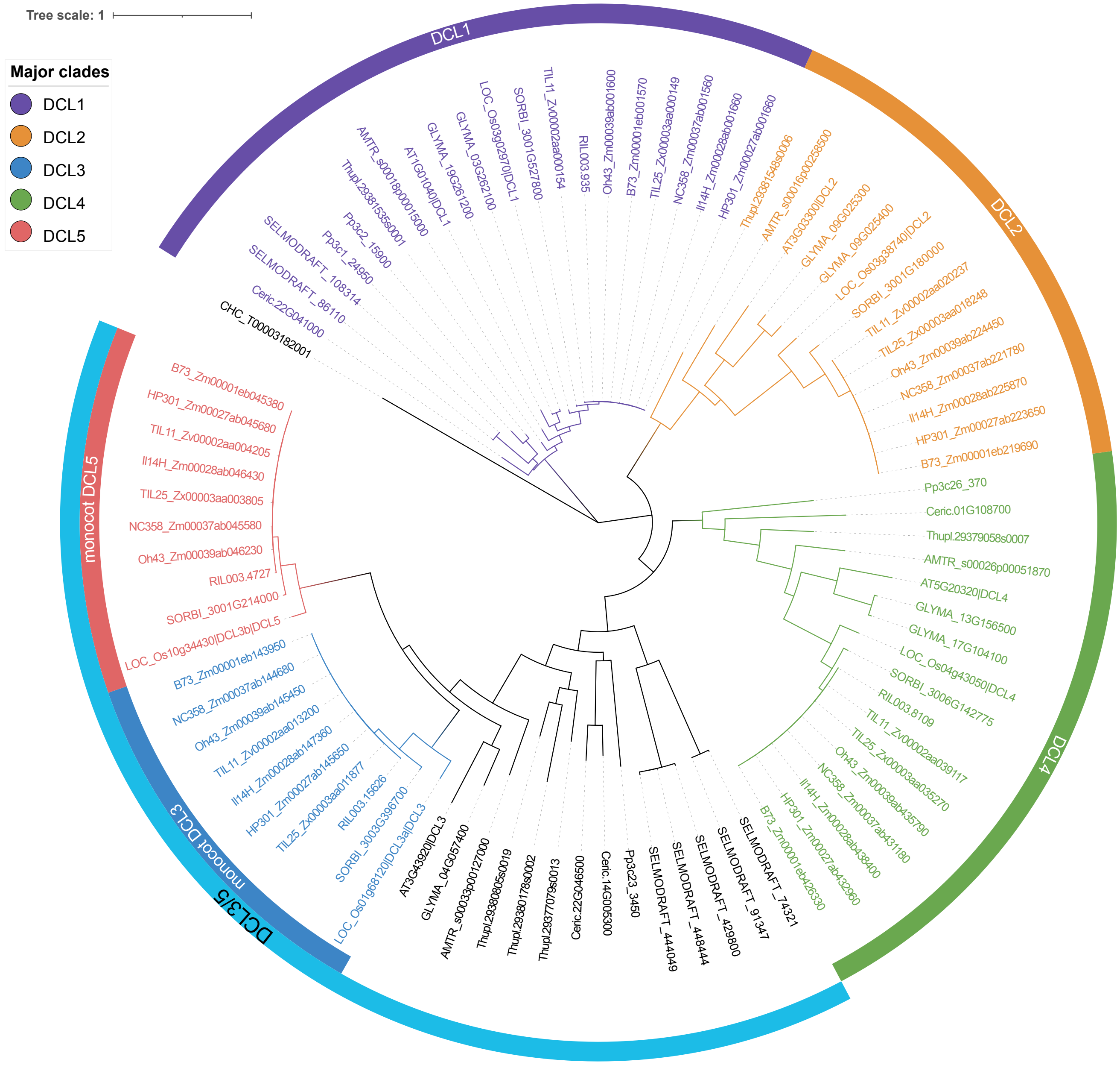

Tree scale: 1
Major clades
DCL1
DCL2
DCL3
DCL4
DCL5
DCL1
DCL2
DCL4
DCL3/5
monocot DCL3
monocot DCL5
TIL11_Zv00002aa000154
RIL003.935
Oh43_Zm00039ab001600
B73_Zm00001eb001570
TIL25_Zx00003aa000149
NC358_Zm00037ab001560
Il14H_Zm00028ab001660
HP301_Zm00027ab001660
SORBI_3001G527800
LOC_Os03g02970|DCL1
GLYMA_03G262100
GLYMA_19G261200
AT1G01040|DCL1
AMTR_s00018p00015000
Thupl.29381535s0001
Pp3c2_15900
Pp3c1_24950
SELMODRAFT_108314
SELMODRAFT_86110
Ceric.22G041000
CHC_T00003182001
Thupl.29381548s0006
AMTR_s00016p00258500
AT3G03300|DCL2
GLYMA_09G025300
GLYMA_09G025400
LOC_Os03g38740|DCL2
SORBI_3001G180000
TIL11_Zv00002aa020237
TIL25_Zx00003aa018248
Oh43_Zm00039ab224450
NC358_Zm00037ab221780
Il14H_Zm00028ab225870
HP301_Zm00027ab223650
B73_Zm00001eb219690
Pp3c26_370
Ceric.01G108700
Thupl.29379058s0007
AMTR_s00026p00051870
AT5G20320|DCL4
GLYMA_13G156500
GLYMA_17G104100
LOC_Os04g43050|DCL4
SORBI_3006G142775
RIL003.8109
TIL11_Zv00002aa039117
TIL25_Zx00003aa035270
Oh43_Zm00039ab435790
NC358_Zm00037ab431180
Il14H_Zm00028ab438400
HP301_Zm00027ab432960
B73_Zm00001eb426330
SELMODRAFT_74321
SELMODRAFT_91347
SELMODRAFT_429800
SELMODRAFT_448444
SELMODRAFT_444049
Pp3c23_3450
Ceric.14G005300
Ceric.22G046500
Thupl.29377079s0013
Thupl.29380178s0002
Thupl.29380805s0019
AMTR_s00033p00127000
GLYMA_04G057400
AT3G43920|DCL3
LOC_Os01g68120|DCL3a|DCL3
SORBI_3003G396700
RIL003.15626
TIL25_Zx00003aa011877
HP301_Zm00027ab145650
Il14H_Zm00028ab147360
TIL11_Zv00002aa013200
Oh43_Zm00039ab145450
NC358_Zm00037ab144680
B73_Zm00001eb143950
LOC_Os10g34430|DCL3b|DCL5
SORBI_3001G214000
RIL003.4727
Oh43_Zm00039ab046230
NC358_Zm00037ab045580
TIL25_Zx00003aa003805
Il14H_Zm00028ab046430
TIL11_Zv00002aa004205
HP301_Zm00027ab045680
B73_Zm00001eb045380


Figure S1D

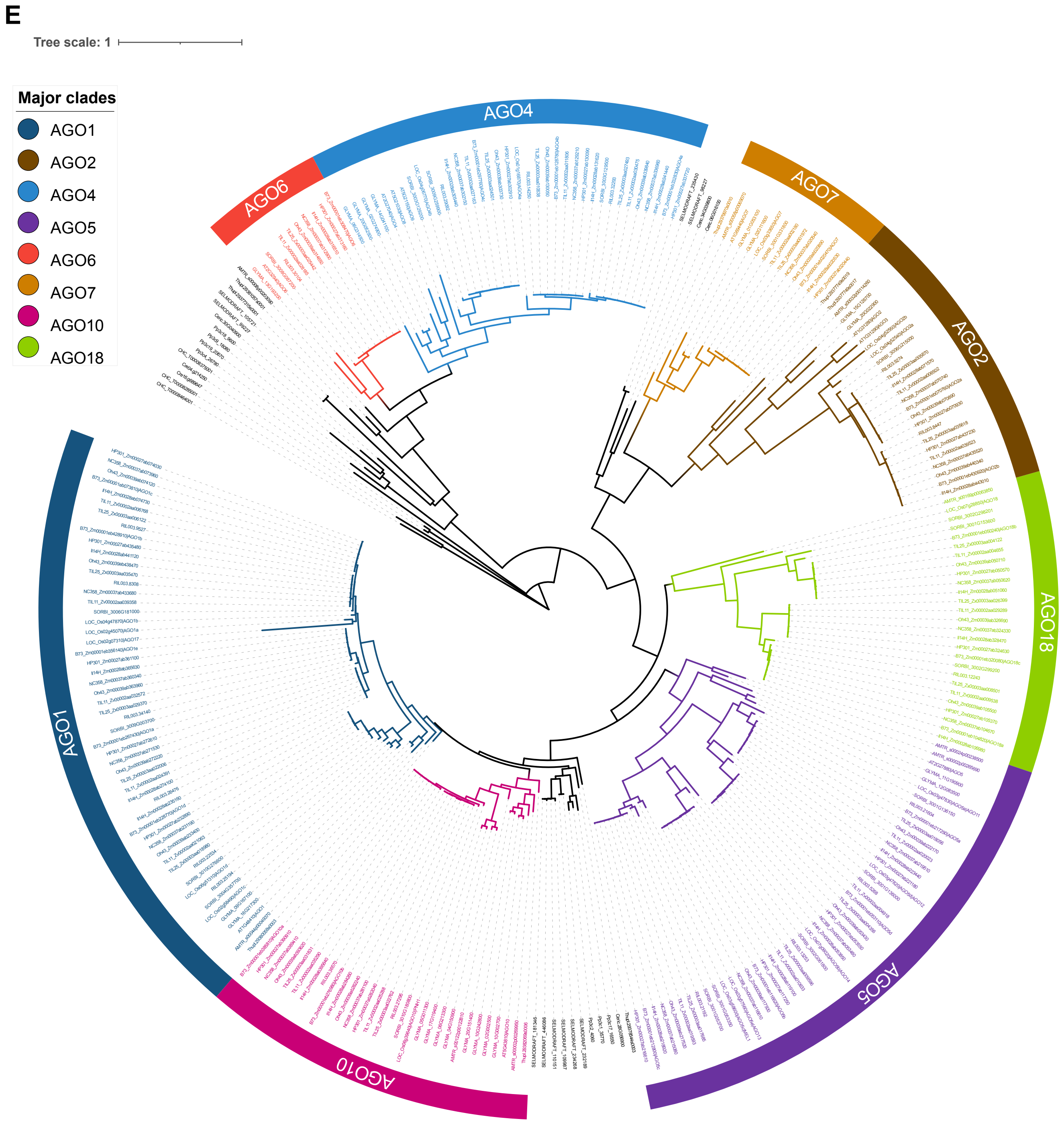
E
Tree scale: 1
Major clades
AGO1
AGO2
AGO4
AGO5
AGO6
AGO7
AGO10
AGO18
AGO4
AGO7
AGO2
AGO18
AGO5
AGO10
AGO1
AGO6

### Figure S1E

F

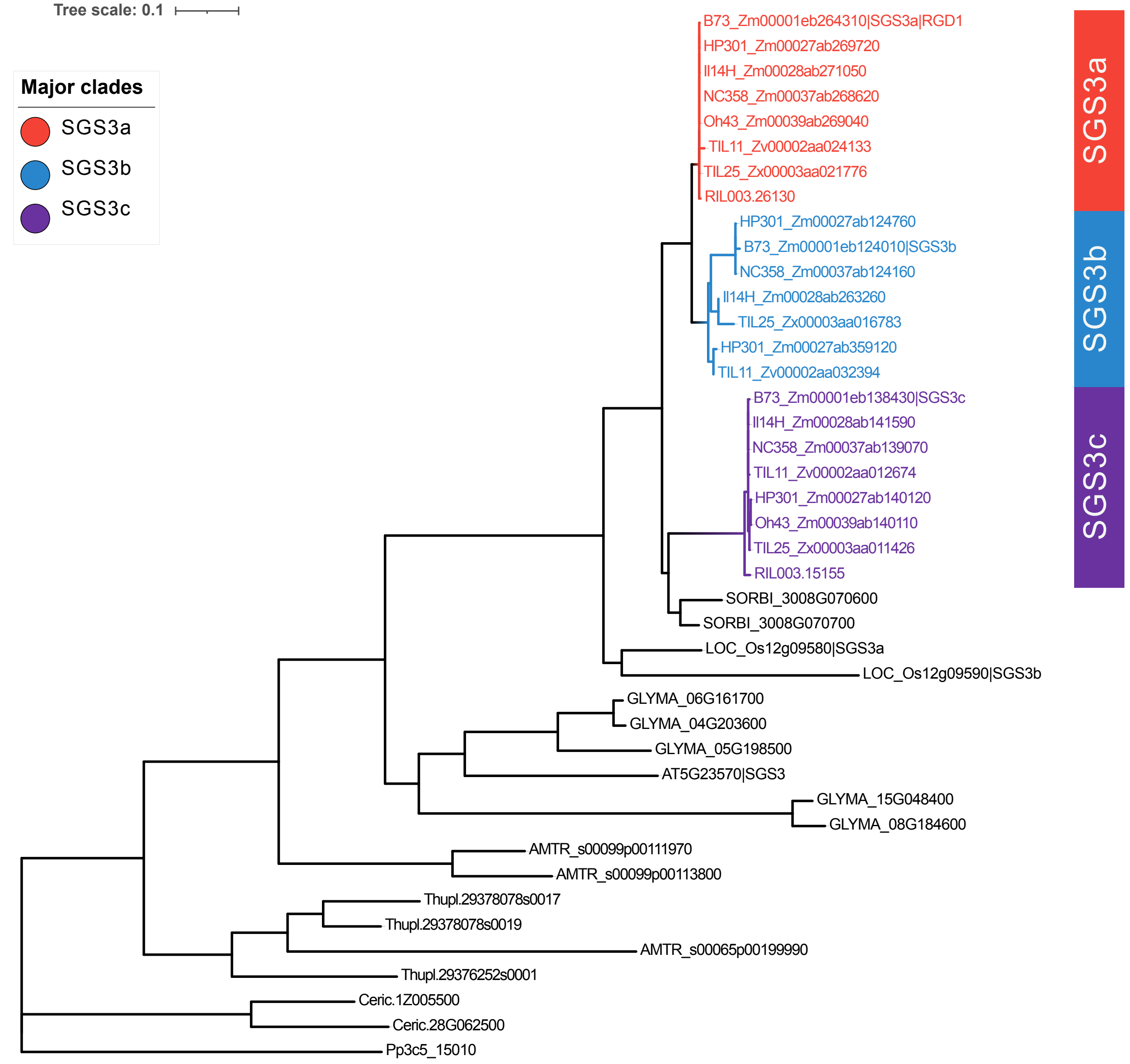

Tree scale: 0.1
Major clades
SGS3a
SGS3b
SGS3c
B73_Zm00001eb264310|SGS3a|RGD1
HP301_Zm00027ab269720
Il14H_Zm00028ab271050
NC358_Zm00037ab268620
Oh43_Zm00039ab269040
TIL11_Zv00002aa024133
TIL25_Zx00003aa021776
RIL003.26130
HP301_Zm00027ab124760
B73_Zm00001eb124010|SGS3b
NC358_Zm00037ab124160
Il14H_Zm00028ab263260
TIL25_Zx00003aa016783
HP301_Zm00027ab359120
TIL11_Zv00002aa032394
B73_Zm00001eb138430|SGS3c
Il14H_Zm00028ab141590
NC358_Zm00037ab139070
TIL11_Zv00002aa012674
HP301_Zm00027ab140120
Oh43_Zm00039ab140110
TIL25_Zx00003aa011426
RIL003.15155
SORBI_3008G070600
SORBI_3008G070700
LOC_Os12g09580|SGS3a
LOC_Os12g09590|SGS3b
GLYMA_06G161700
GLYMA_04G203600
GLYMA_05G198500
AT5G23570|SGS3
GLYMA_15G048400
GLYMA_08G184600
AMTR_s00099p00111970
AMTR_s00099p00113800
Thupl.29378078s0017
Thupl.29378078s0019
AMTR_s00065p00199990
Thupl.29376252s0001
Ceric.1Z005500
Ceric.28G062500
Pp3c5_15010
SGS3a
SGS3b
SGS3c


### Figure S1F

G

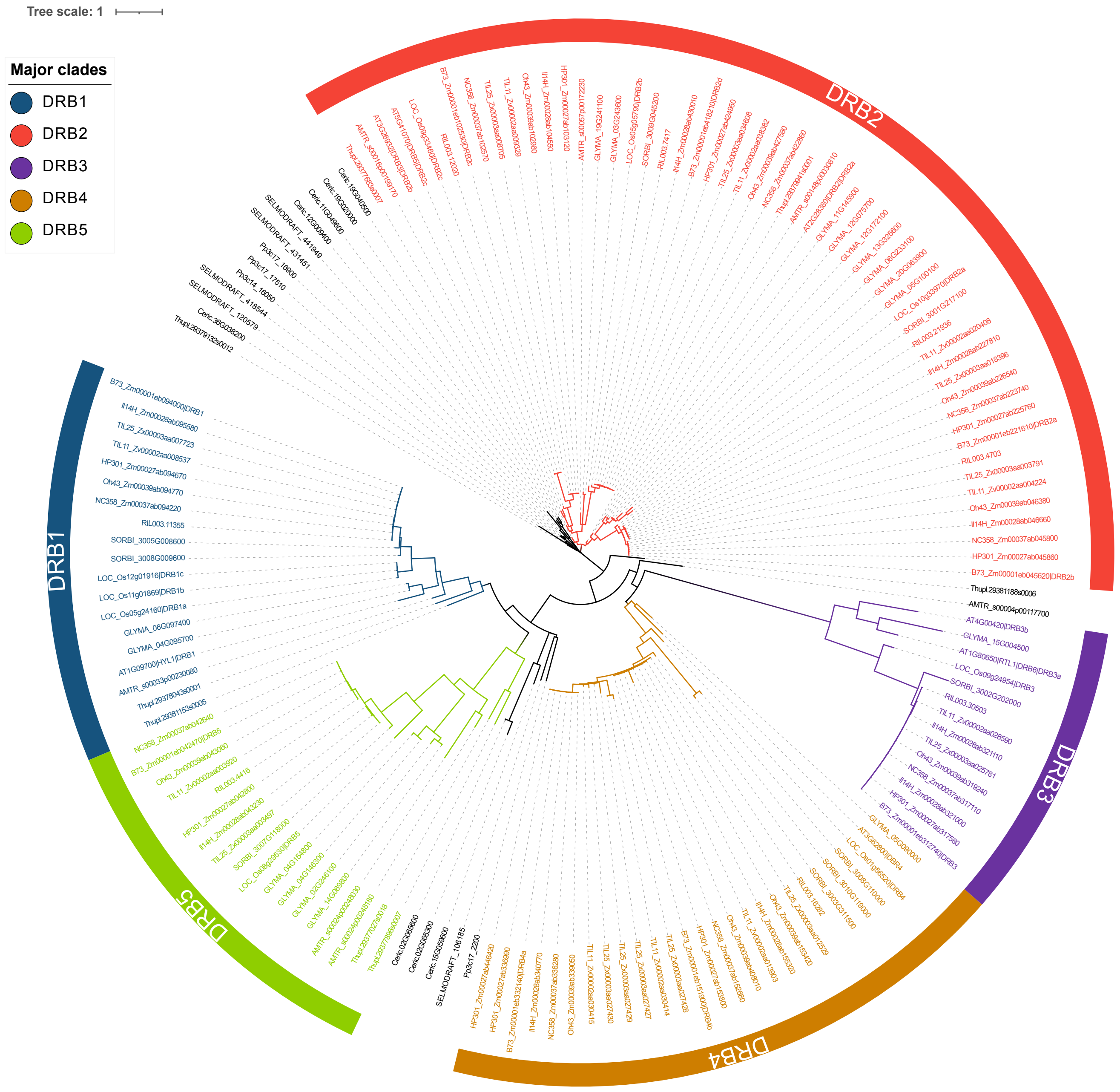
Tree scale: 1
Major clades
DRB1
DRB2
DRB3
DRB4
DRB5
DRB1
DRB2
DRB3
DRB4
DRB5

### Figure S1G

**H**

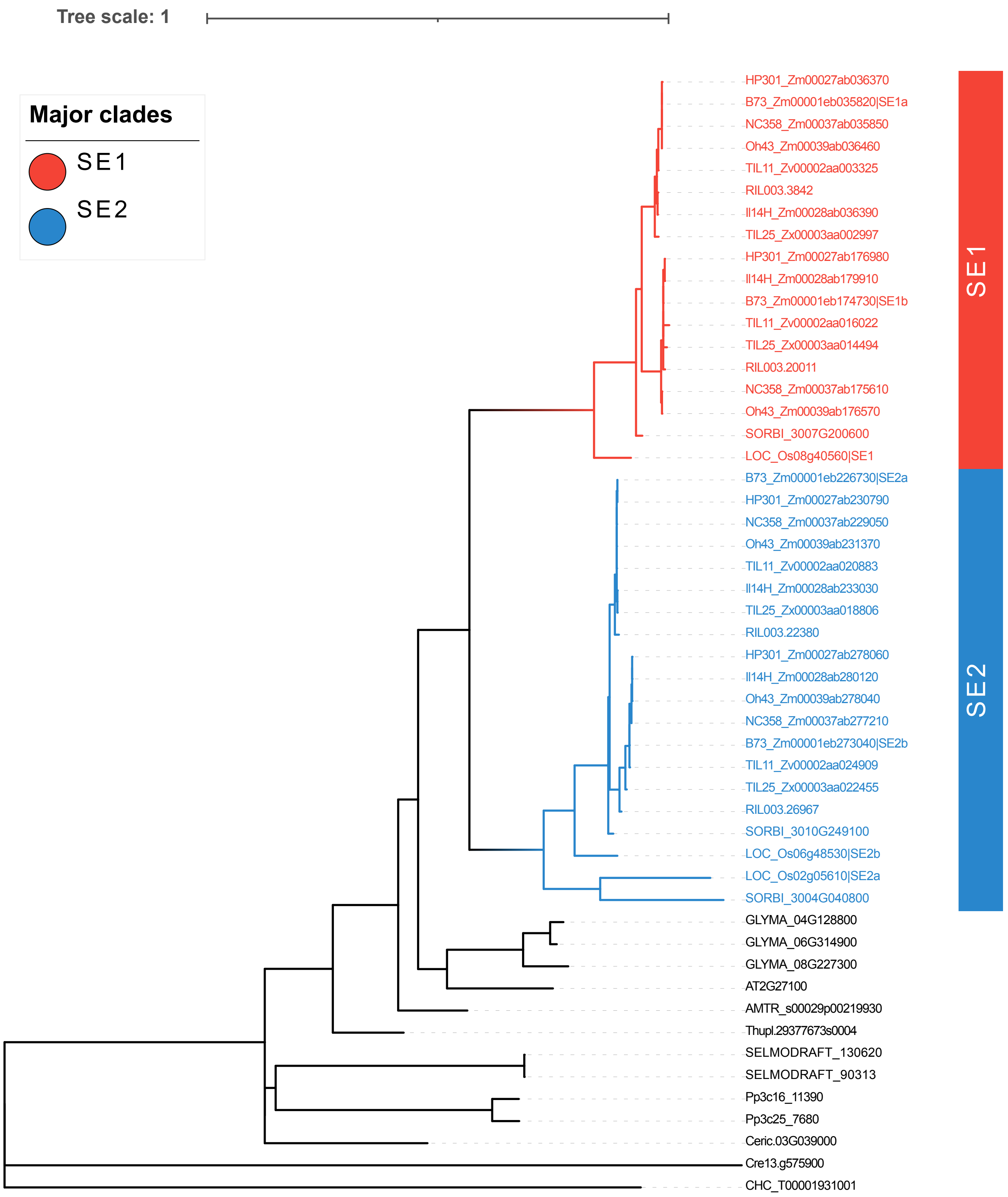

Tree scale: 1
Major clades
SE1
SE2
HP301_Zm00027ab036370
B73_Zm00001eb035820|SE1a
NC358_Zm00037ab035850
Oh43_Zm00039ab036460
TIL11_Zv00002aa003325
RIL003.3842
Il14H_Zm00028ab036390
TIL25_Zx00003aa002997
HP301_Zm00027ab176980
Il14H_Zm00028ab179910
B73_Zm00001eb174730|SE1b
TIL11_Zv00002aa016022
TIL25_Zx00003aa014494
RIL003.20011
NC358_Zm00037ab175610
Oh43_Zm00039ab176570
SORBI_3007G200600
LOC_Os08g40560|SE1
B73_Zm00001eb226730|SE2a
HP301_Zm00027ab230790
NC358_Zm00037ab229050
Oh43_Zm00039ab231370
TIL11_Zv00002aa020883
Il14H_Zm00028ab233030
TIL25_Zx00003aa018806
RIL003.22380
HP301_Zm00027ab278060
Il14H_Zm00028ab280120
Oh43_Zm00039ab278040
NC358_Zm00037ab277210
B73_Zm00001eb273040|SE2b
TIL11_Zv00002aa024909
TIL25_Zx00003aa022455
RIL003.26967
SORBI_3010G249100
LOC_Os06g48530|SE2b
LOC_Os02g05610|SE2a
SORBI_3004G040800
GLYMA_04G128800
GLYMA_06G314900
GLYMA_08G227300
AT2G27100
AMTR_s00029p00219930
Thupl.29377673s0004
SELMODRAFT_130620
SELMODRAFT_90313
Pp3c16_11390
Pp3c25_7680
Ceric.03G039000
Cre13.g575900
CHC_T00001931001
SE1
SE2


### Figure S1H

I

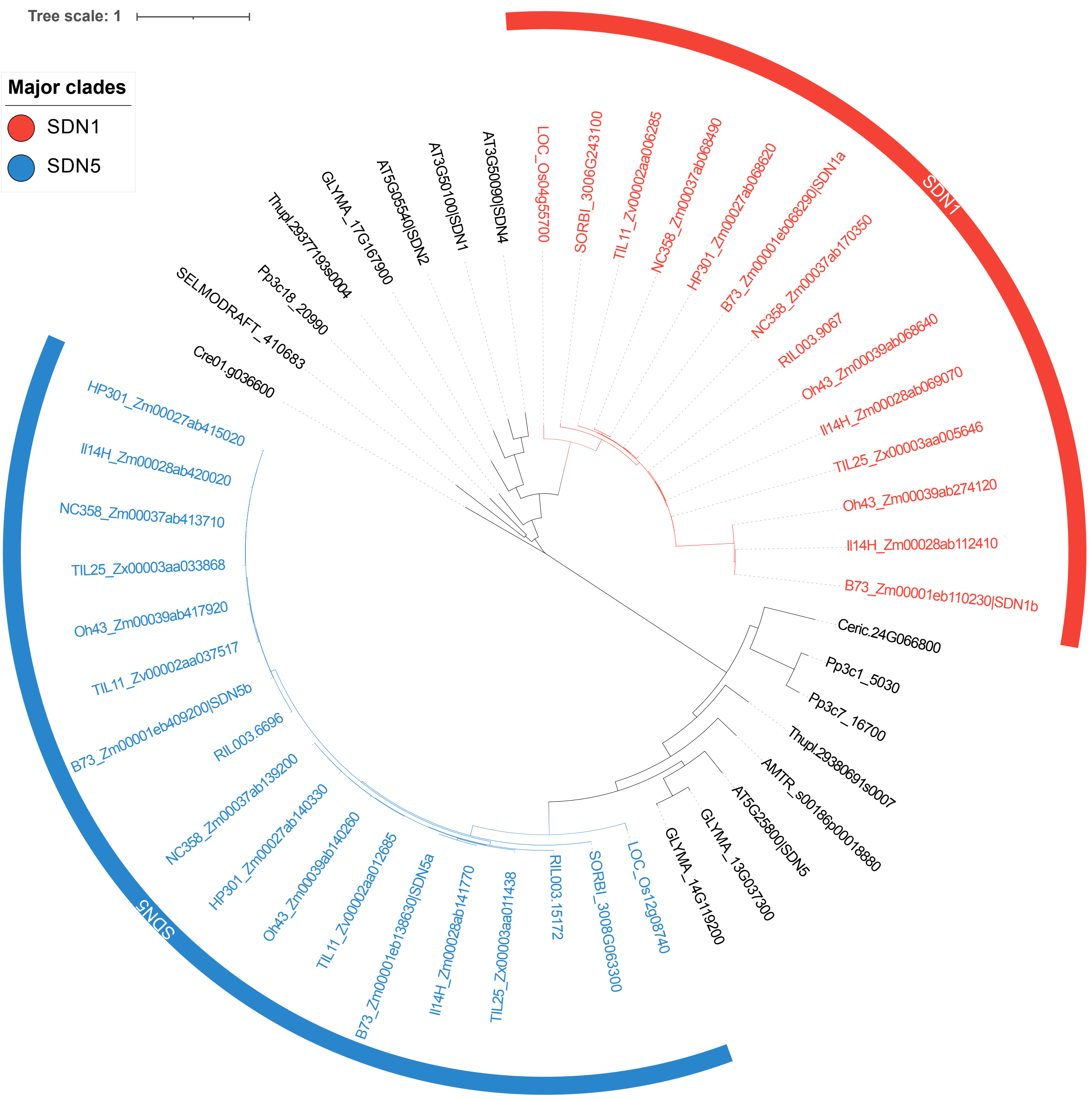

Tree scale: 1
Major clades
SDN1
SDN5
LOC_Os04g55700
SORBI_3006G243100
TIL11_Zv00002aa006285
NC358_Zm00037ab068490
HP301_Zm00027ab068620
B73_Zm00001eb068290|SDN1a
NC358_Zm00037ab170350
RIL003.9067
Oh43_Zm00039ab068640
Il14H_Zm00028ab069070
TIL25_Zx00003aa005646
Oh43_Zm00039ab274120
Il14H_Zm00028ab112410
B73_Zm00001eb110230|SDN1b
Ceric.24G066800
Pp3c1_5030
Pp3c7_16700
Thupl.29380691s0007
AMTR_s00186p00018880
AT5G25800|SDN5
GLYMA_13G037300
GLYMA_14G119200
LOC_Os12g08740
SORBI_3008G063300
RIL003.15172
TIL25_Zx00003aa011438
Il14H_Zm00028ab141770
B73_Zm00001eb138630|SDN5a
TIL11_Zv00002aa012685
Oh43_Zm00039ab140260
HP301_Zm00027ab140330
NC358_Zm00037ab139200
RIL003.6696
B73_Zm00001eb409200|SDN5b
TIL11_Zv00002aa037517
Oh43_Zm00039ab417920
TIL25_Zx00003aa033868
NC358_Zm00037ab413710
Il14H_Zm00028ab420020
HP301_Zm00027ab415020
Cre01.g036600
SELMODRAFT_410683
Pp3c18_20990
Thupl.29377193s0004
GLYMA_17G167900
AT5G05540|SDN2
AT3G50100|SDN1
AT3G50090|SDN4
SDN1
SDN5


### Figure S1I

**J**

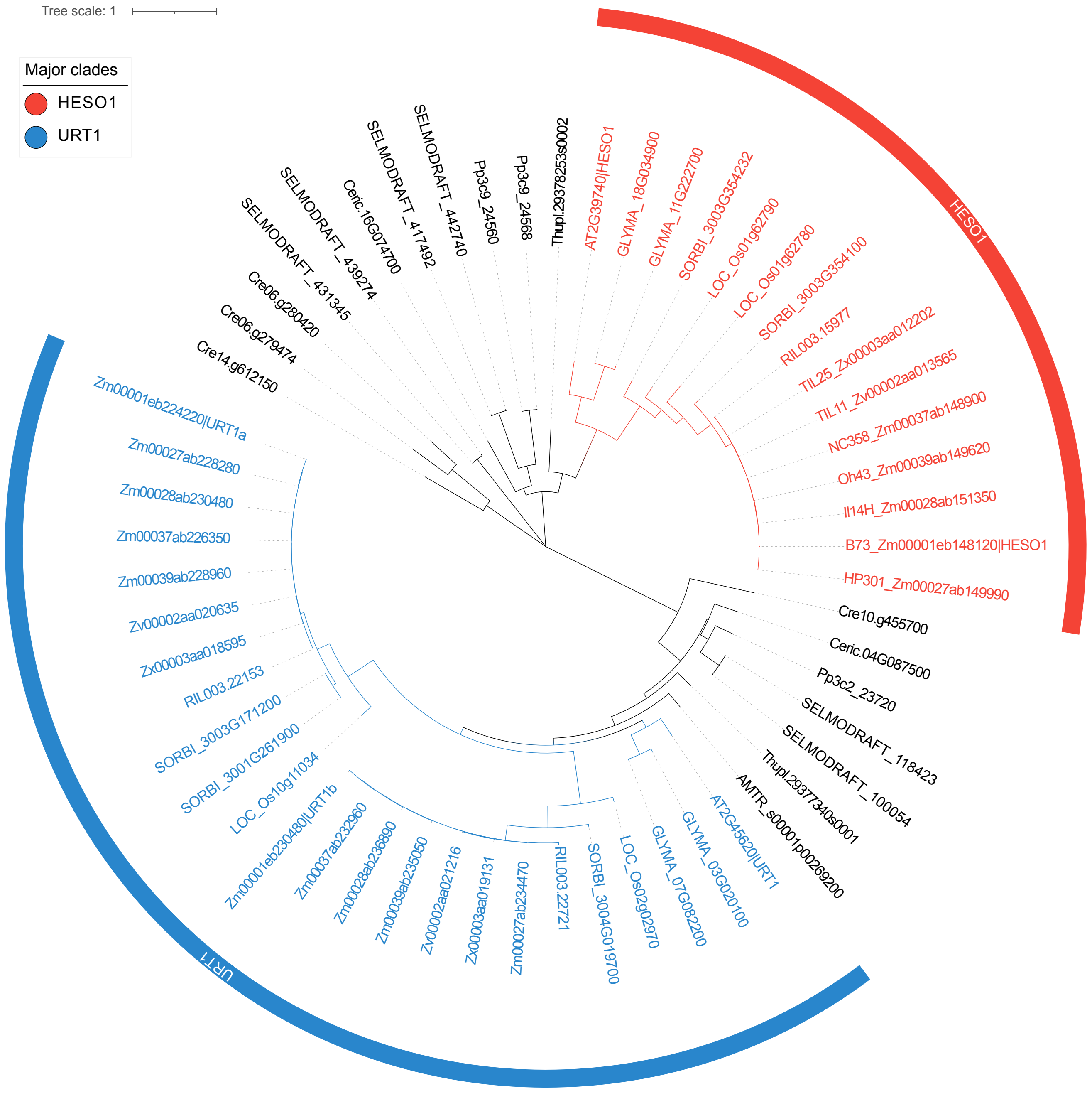
Tree scale: 1
Major clades
HESO1
URT1
AT2G39740|HESO1
GLYMA_18G034900
GLYMA_11G222700
SORBI_3003G354232
LOC_Os01g62790
LOC_Os01g62780
SORBI_3003G354100
RIL003.15977
TIL25_Zx00003aa012202
TIL11_Zv00002aa013565
NC358_Zm00037ab148900
Oh43_Zm00039ab149620
Il14H_Zm00028ab151350
B73_Zm00001eb148120|HESO1
HP301_Zm00027ab149990
Cre10.g455700
Ceric.04G087500
Pp3c2_23720
SELMODRAFT_118423
SELMODRAFT_100054
Thupl.29377340s0001
AMTR_s00001p00269200
AT2G45620|URT1
GLYMA_03G020100
GLYMA_07G082200
LOC_Os02g02970
SORBI_3004G019700
RIL003.22721
Zm00027ab234470
Zx00003aa019131
Zv00002aa021216
Zm00039ab235050
Zm00028ab236890
Zm00037ab232960
Zm00001eb230480|URT1b
LOC_Os10g11034
SORBI_3001G261900
SORBI_3003G171200
RIL003.22153
Zx00003aa018595
Zv00002aa020635
Zm00039ab228960
Zm00037ab226350
Zm00028ab230480
Zm00027ab228280
Zm00001eb224220|URT1a
Cre14.g612150
Cre06.g279474
Cre06.g280420
SELMODRAFT_431345
SELMODRAFT_439274
Ceric.16G074700
SELMODRAFT_417492
SELMODRAFT_442740
Pp3c9_24560
Pp3c9_24568
Thupl.29378253s0002

### Figure S1J

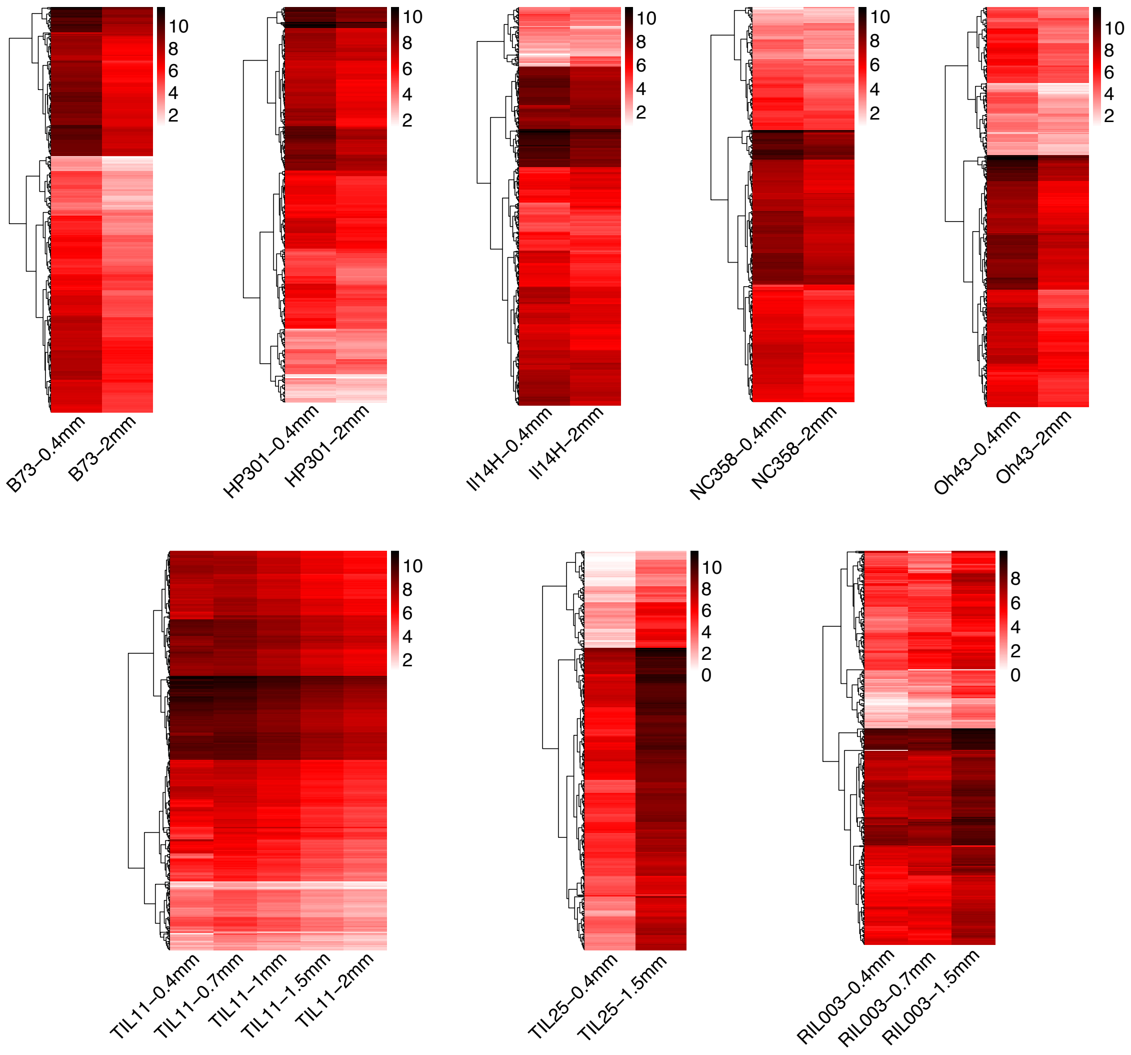
B73–0.4mm
B73–2mm
HP301–0.4mm
HP301–2mm
Il14H–0.4mm
Il14H–2mm
NC358–0.4mm
NC358–2mm
Oh43–0.4mm
Oh43–2mm
TIL11–0.4mm
TIL11–0.7mm
TIL11–1mm
TIL11–1.5mm
TIL11–2mm
TIL25–0.4mm
TIL25–1.5mm
RIL003–0.4mm
RIL003–0.7mm
RIL003–1.5mm

Figure S2

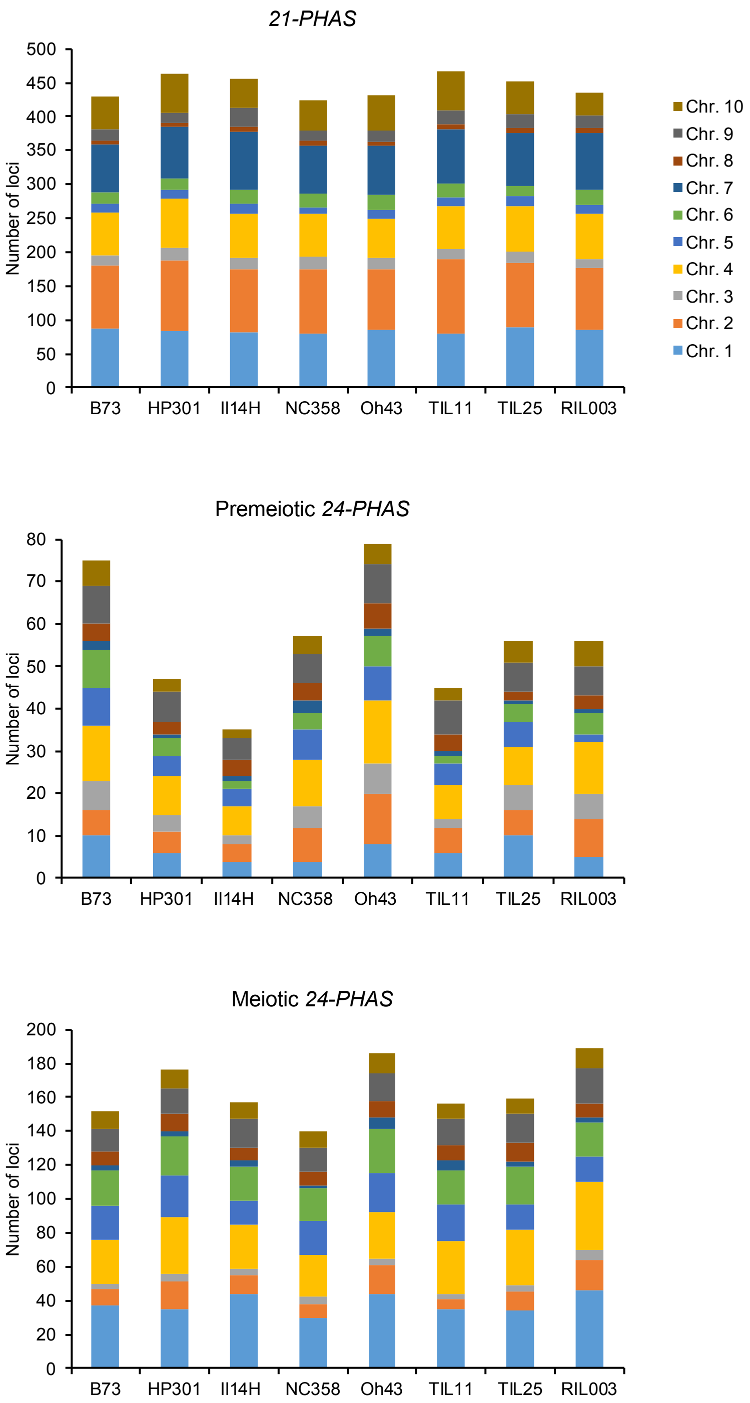

21-PHAS
Number of loci
0
50
100
150
200
250
300
350
400
450
500
B73
HP301
Il14H
NC358
Oh43
TIL11
TIL25
RIL003
Chr. 10
Chr. 9
Chr. 8
Chr. 7
Chr. 6
Chr. 5
Chr. 4
Chr. 3
Chr. 2
Chr. 1
Premeiotic 24-PHAS
Number of loci
0
10
20
30
40
50
60
70
80
B73
HP301
Il14H
NC358
Oh43
TIL11
TIL25
RIL003
Meiotic 24-PHAS
Number of loci
0
20
40
60
80
100
120
140
160
180
200
B73
HP301
Il14H
NC358
Oh43
TIL11
TIL25
RIL003


### Figure S3

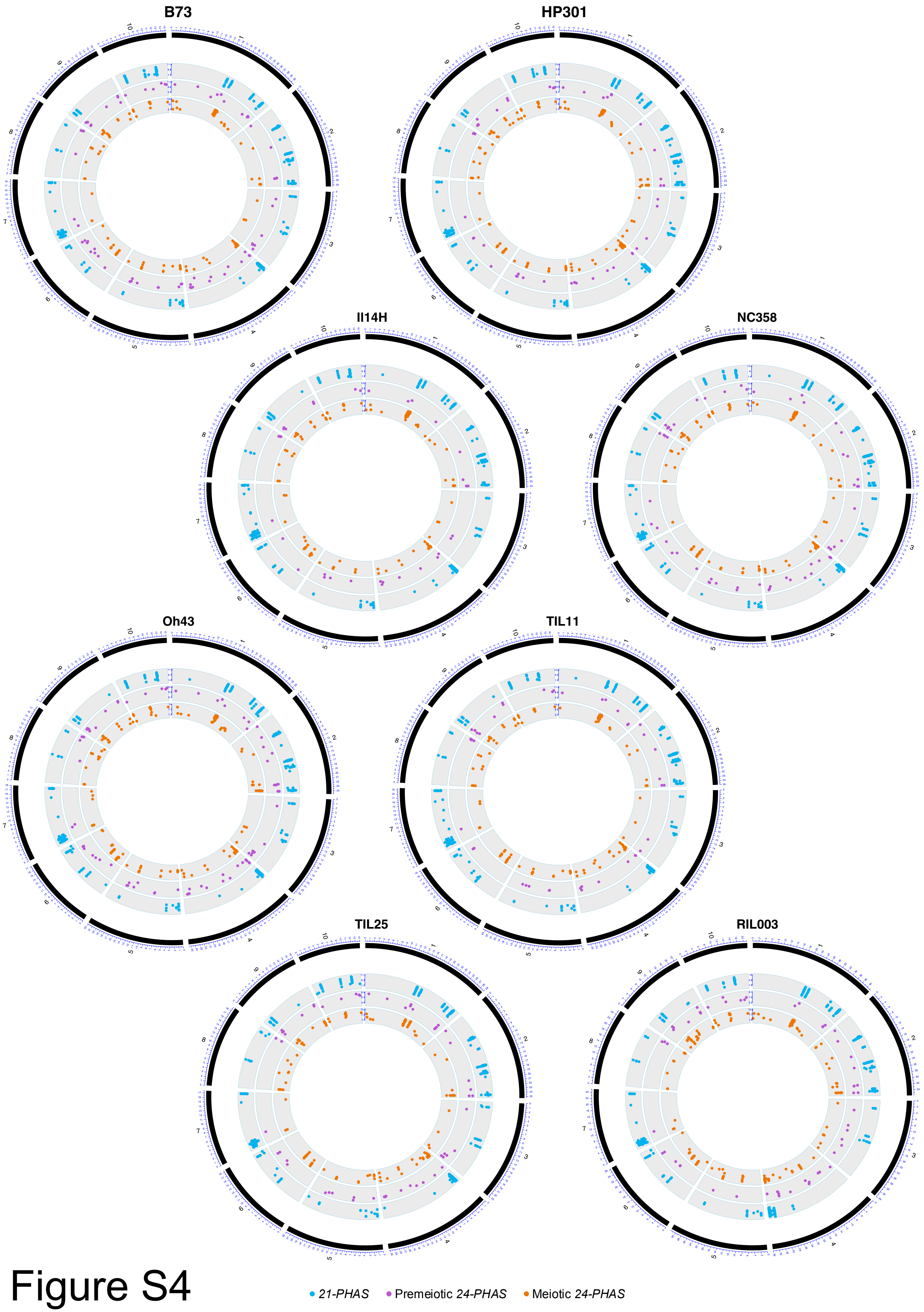
B73
HP301
Il14H
NC358
Oh43
TIL11
TIL25
RIL003
21-PHAS
Premeiotic 24-PHAS
Meiotic 24-PHAS

Figure S4

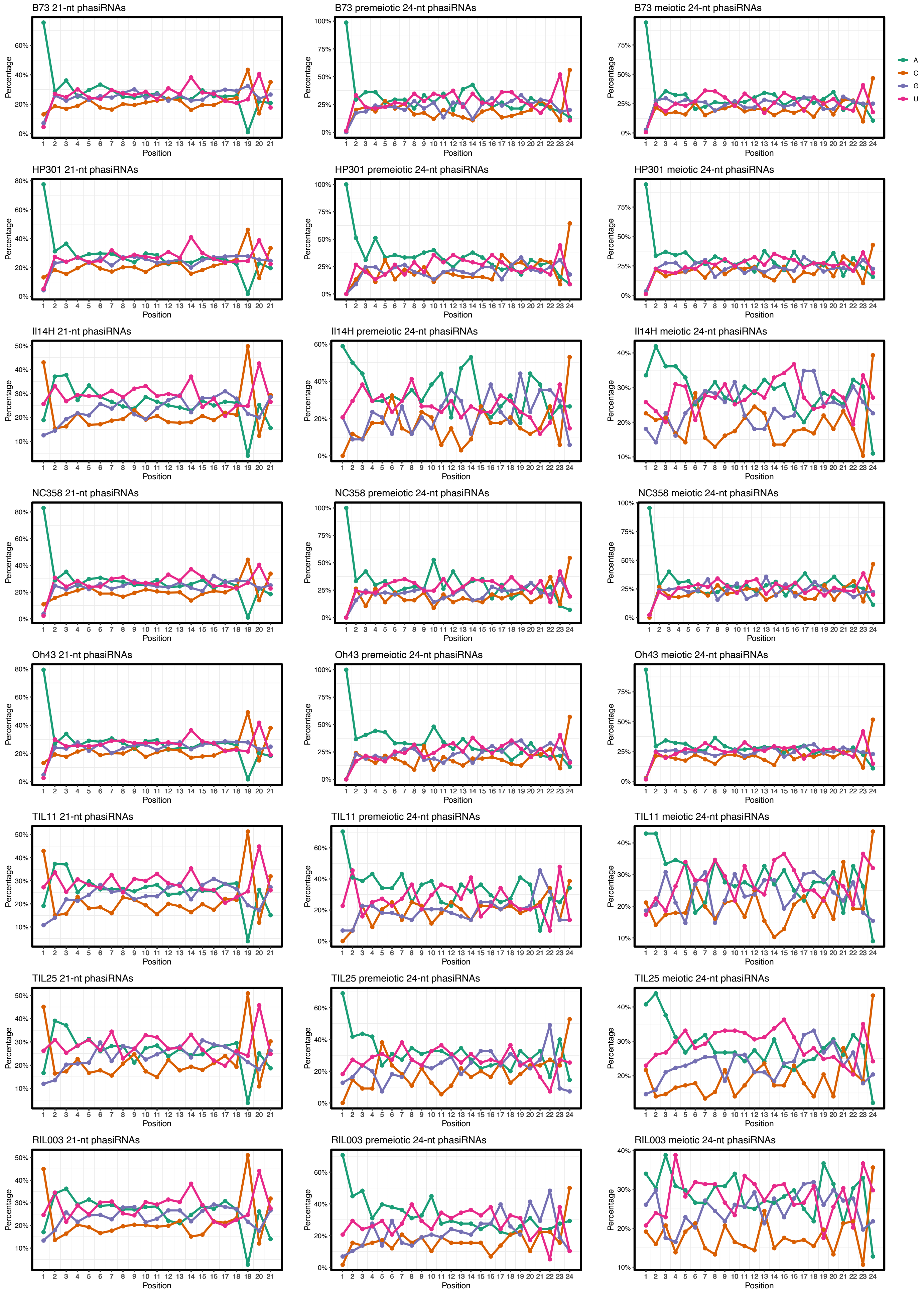
B73 21-nt phasiRNAs
B73 premeiotic 24-nt phasiRNAs
B73 meiotic 24-nt phasiRNAs
HP301 21-nt phasiRNAs
HP301 premeiotic 24-nt phasiRNAs
HP301 meiotic 24-nt phasiRNAs
Il14H 21-nt phasiRNAs
Il14H premeiotic 24-nt phasiRNAs
Il14H meiotic 24-nt phasiRNAs
NC358 21-nt phasiRNAs
NC358 premeiotic 24-nt phasiRNAs
NC358 meiotic 24-nt phasiRNAs
Oh43 21-nt phasiRNAs
Oh43 premeiotic 24-nt phasiRNAs
Oh43 meiotic 24-nt phasiRNAs
TIL11 21-nt phasiRNAs
TIL11 premeiotic 24-nt phasiRNAs
TIL11 meiotic 24-nt phasiRNAs
TIL25 21-nt phasiRNAs
TIL25 premeiotic 24-nt phasiRNAs
TIL25 meiotic 24-nt phasiRNAs
RIL003 21-nt phasiRNAs
RIL003 premeiotic 24-nt phasiRNAs
RIL003 meiotic 24-nt phasiRNAs
Percentage
Position
A
C
G
U

### Figure S5

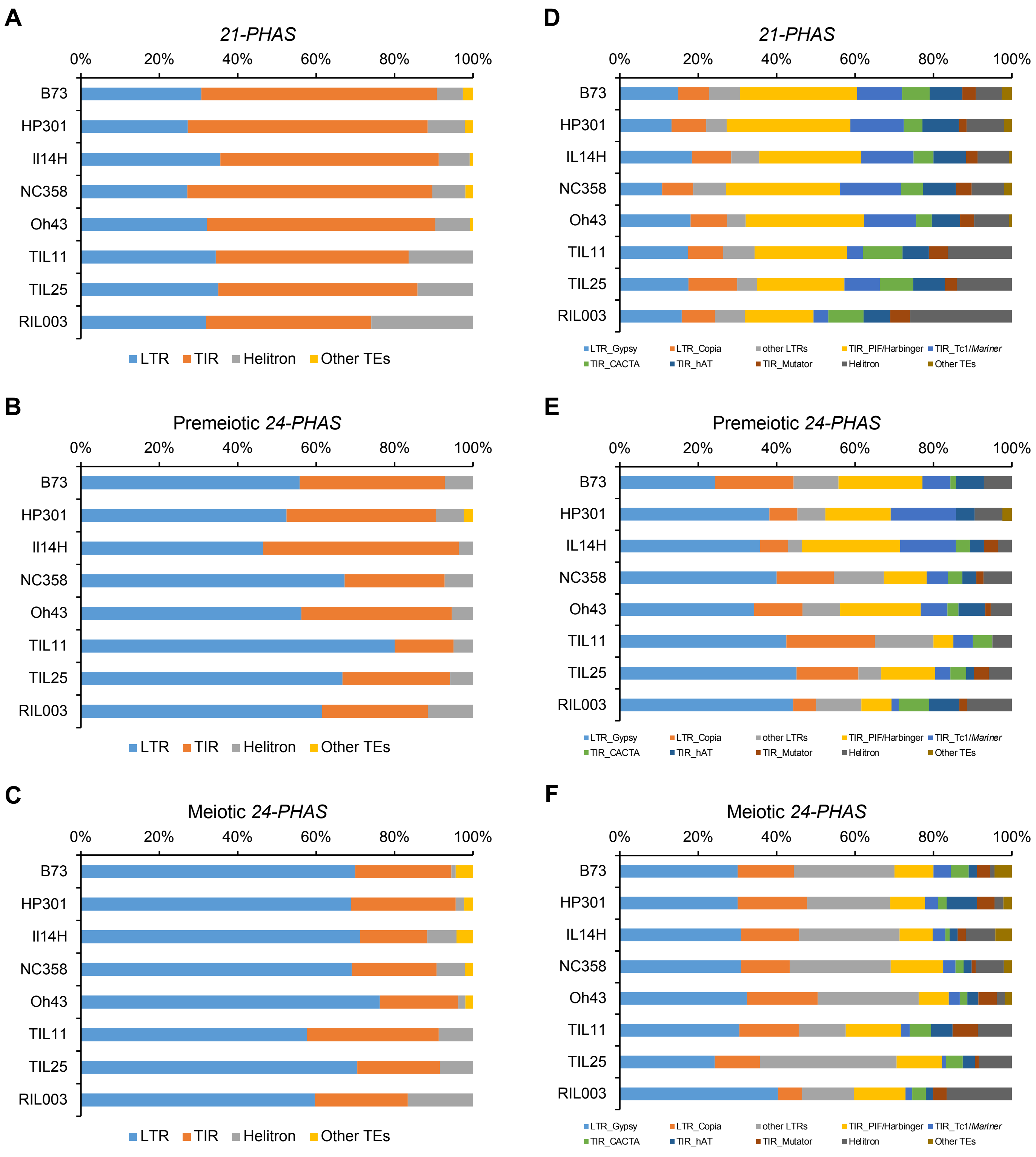
A
21-PHAS
0% 20% 40% 60% 80% 100%
B73
HP301
Il14H
NC358
Oh43
TIL11
TIL25
RIL003
LTR TIR Helitron Other TEs
B
Premeiotic 24-PHAS
0% 20% 40% 60% 80% 100%
B73
HP301
Il14H
NC358
Oh43
TIL11
TIL25
RIL003
LTR TIR Helitron Other TEs
C
Meiotic 24-PHAS
0% 20% 40% 60% 80% 100%
B73
HP301
Il14H
NC358
Oh43
TIL11
TIL25
RIL003
LTR TIR Helitron Other TEs
D
21-PHAS
0% 20% 40% 60% 80% 100%
B73
HP301
IL14H
NC358
Oh43
TIL11
TIL25
RIL003
LTR_Gypsy LTR_Copia other LTRs TIR_PIF/Harbinger TIR_Tc1/Mariner
TIR_CACTA TIR_hAT TIR_Mutator Helitron Other TEs
E
Premeiotic 24-PHAS
0% 20% 40% 60% 80% 100%
B73
HP301
IL14H
NC358
Oh43
TIL11
TIL25
RIL003
LTR_Gypsy LTR_Copia other LTRs TIR_PIF/Harbinger TIR_Tc1/Mariner
TIR_CACTA TIR_hAT TIR_Mutator Helitron Other TEs
F
Meiotic 24-PHAS
0% 20% 40% 60% 80% 100%
B73
HP301
IL14H
NC358
Oh43
TIL11
TIL25
RIL003
LTR_Gypsy LTR_Copia other LTRs TIR_PIF/Harbinger TIR_Tc1/Mariner
TIR_CACTA TIR_hAT TIR_Mutator Helitron Other TEs

### Figure S6

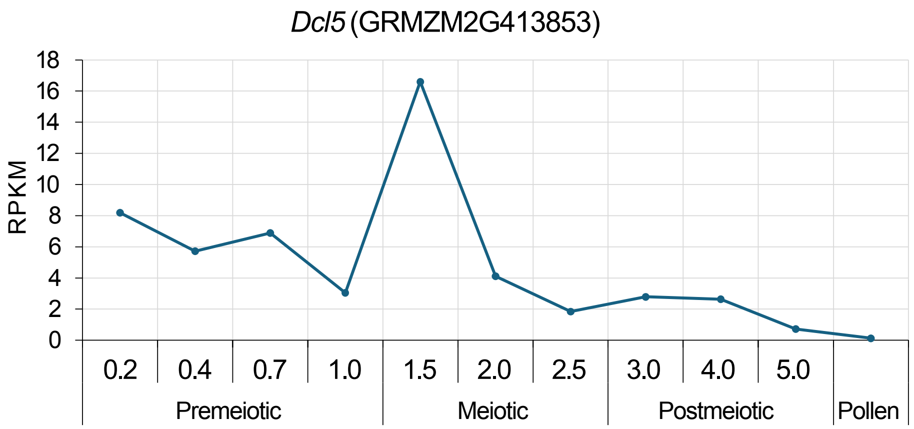

*Dcl5* (GRMZM2G413853)
RPKM
18
16
14
12
10
8
6
4
2
0
0.2
0.4
0.7
1.0
1.5
2.0
2.5
3.0
4.0
5.0
Premeiotic
Meiotic
Postmeiotic
Pollen


### Figure S7

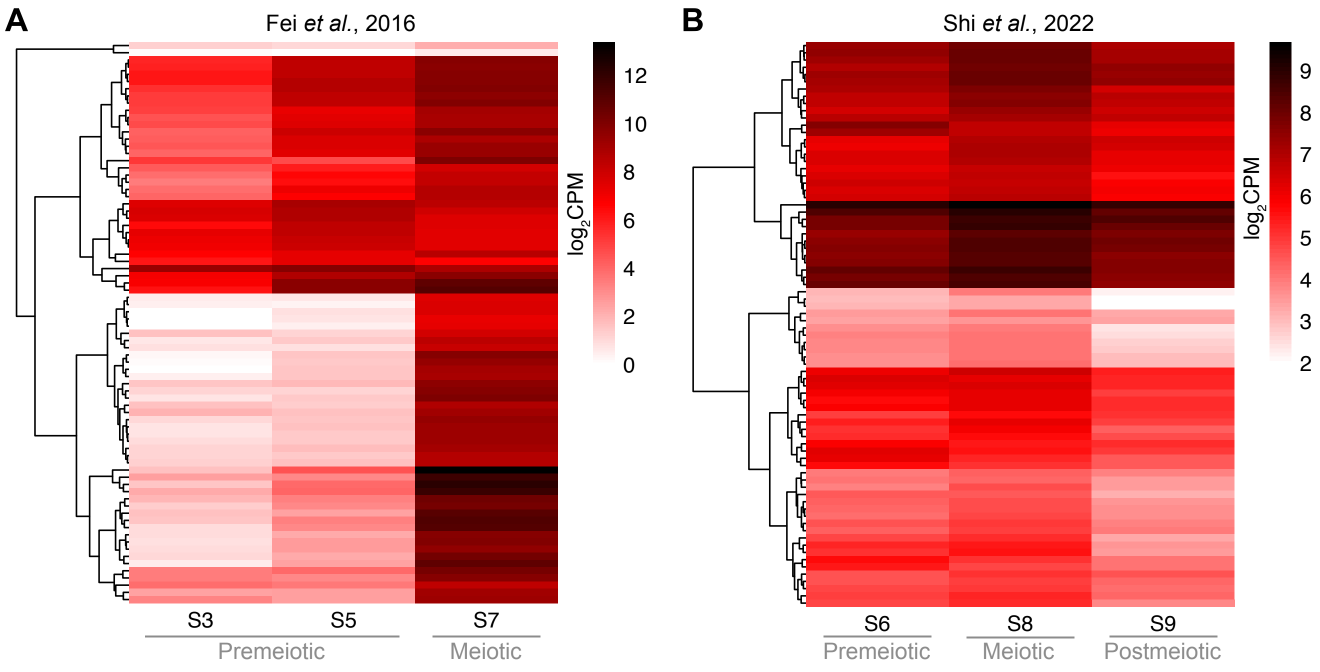
A
Fei *et al.*, 2016
12
10
8
6
4
2
0
$log_2$CPM
S3
S5
S7
Premeiotic
Meiotic
B
Shi *et al.*, 2022
9
8
7
6
5
4
3
2
$log_2$CPM
S6
S8
S9
Premeiotic
Meiotic
Postmeiotic

Figure S8

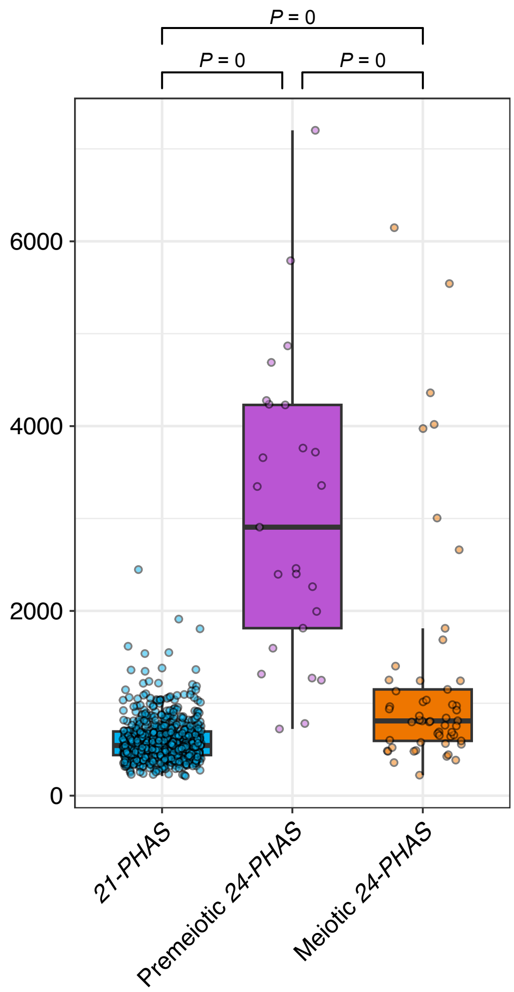

P = 0
P = 0
P = 0
6000
4000
2000
0
21-PHAS
Premeiotic 24-PHAS
Meiotic 24-PHAS


### Figure S9

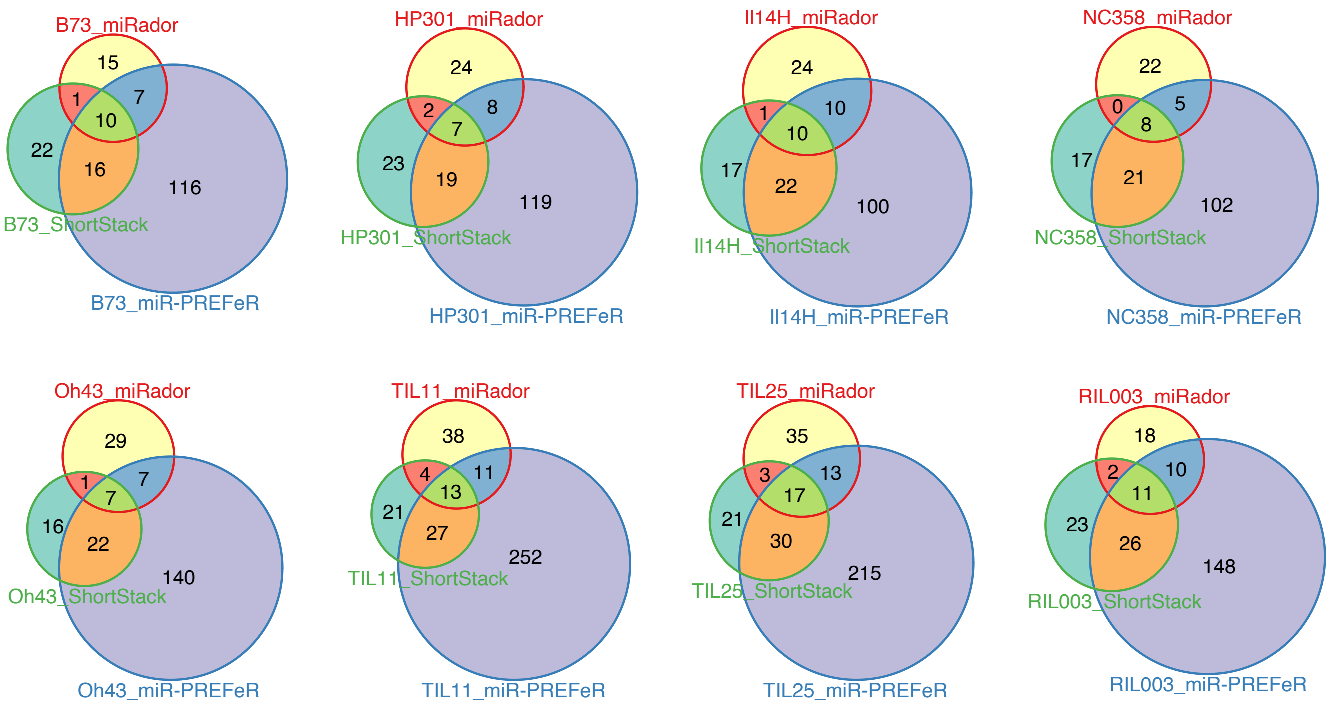
B73_miRador
15
1
7
10
22
16
116
B73_ShortStack
B73_miR-PREFeR
HP301_miRador
24
2
8
7
23
19
119
HP301_ShortStack
HP301_miR-PREFeR
Il14H_miRador
24
1
10
10
17
22
100
Il14H_ShortStack
Il14H_miR-PREFeR
NC358_miRador
22
0
5
8
17
21
102
NC358_ShortStack
NC358_miR-PREFeR
Oh43_miRador
29
1
7
7
16
22
140
Oh43_ShortStack
Oh43_miR-PREFeR
TIL11_miRador
38
4
11
13
21
27
252
TIL11_ShortStack
TIL11_miR-PREFeR
TIL25_miRador
35
3
13
17
21
30
215
TIL25_ShortStack
TIL25_miR-PREFeR
RIL003_miRador
18
2
10
11
23
26
148
RIL003_ShortStack
RIL003_miR-PREFeR

### Figure S10

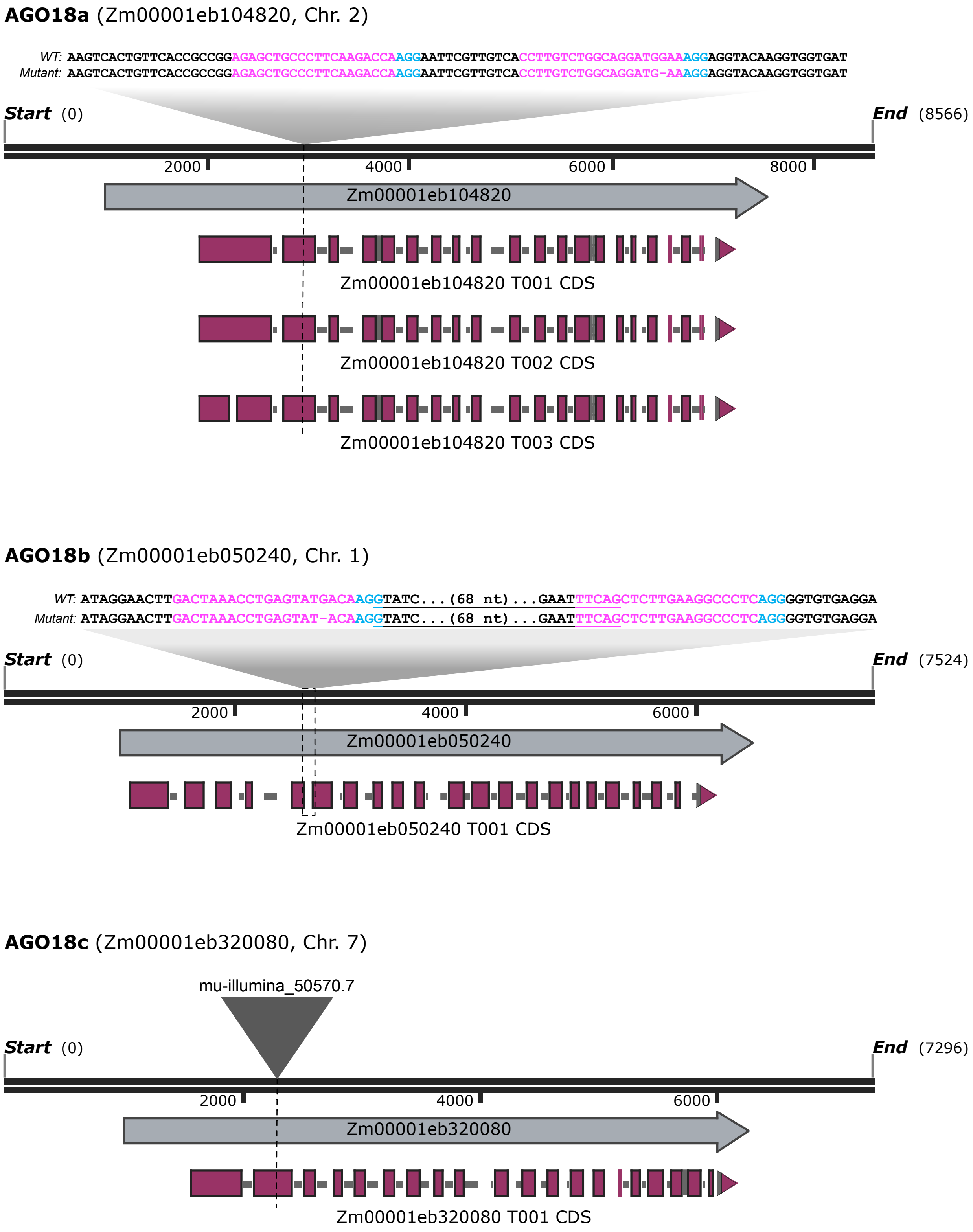
AGO18a (Zm00001eb104820, Chr. 2)
WT: AAGTCACTGTTCACCGCCGGAGAGCTGCCCTTCAAGACCAAGGAATTCGTTGTCACCTTGTCTGGCAGGATGGAAAGGAGGTACAAGGTGGTGAT
Mutant: AAGTCACTGTTCACCGCCGGAGAGCTGCCCTTCAAGACCAAGGAATTCGTTGTCACCTTGTCTGGCAGGATG-AAAGGAGGTACAAGGTGGTGAT
Start (0)
End (8566)
2000
4000
6000
8000
Zm00001eb104820
Zm00001eb104820 T001 CDS
Zm00001eb104820 T002 CDS
Zm00001eb104820 T003 CDS
AGO18b (Zm00001eb050240, Chr. 1)
WT: ATAGGAACTTGACTAAACCTGAGTATGACAAGGTATC...(68 nt)...GAATTTCAGCTCTTGAAGGCCCTCAGGGGTGTGAGGA
Mutant: ATAGGAACTTGACTAAACCTGAGTAT-ACAAGGTATC...(68 nt)...GAATTTCAGCTCTTGAAGGCCCTCAGGGGTGTGAGGA
Start (0)
End (7524)
2000
4000
6000
Zm00001eb050240
Zm00001eb050240 T001 CDS
AGO18c (Zm00001eb320080, Chr. 7)
mu-illumina_50570.7
Start (0)
End (7296)
2000
4000
6000
Zm00001eb320080
Zm00001eb320080 T001 CDS

### Figure S11

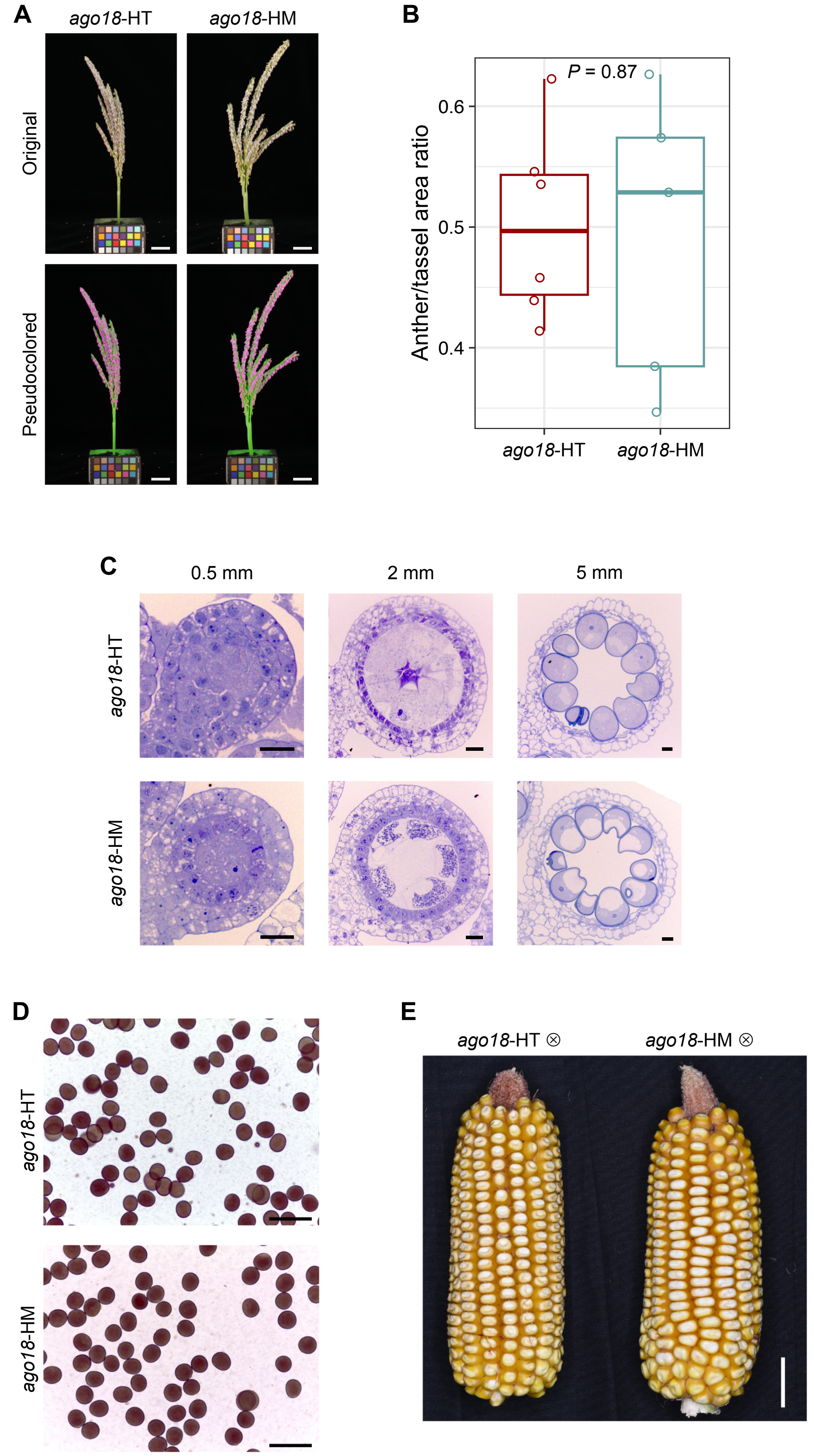
A
ago18-HT
ago18-HM
Original
Pseudocolored
B
P = 0.87
0.6
0.5
0.4
Anther/tassel area ratio
ago18-HT
ago18-HM
C
0.5 mm
2 mm
5 mm
ago18-HT
ago18-HM
D
ago18-HT
ago18-HM
E
ago18-HT ⊗
ago18-HM ⊗

Figure S12

**A**

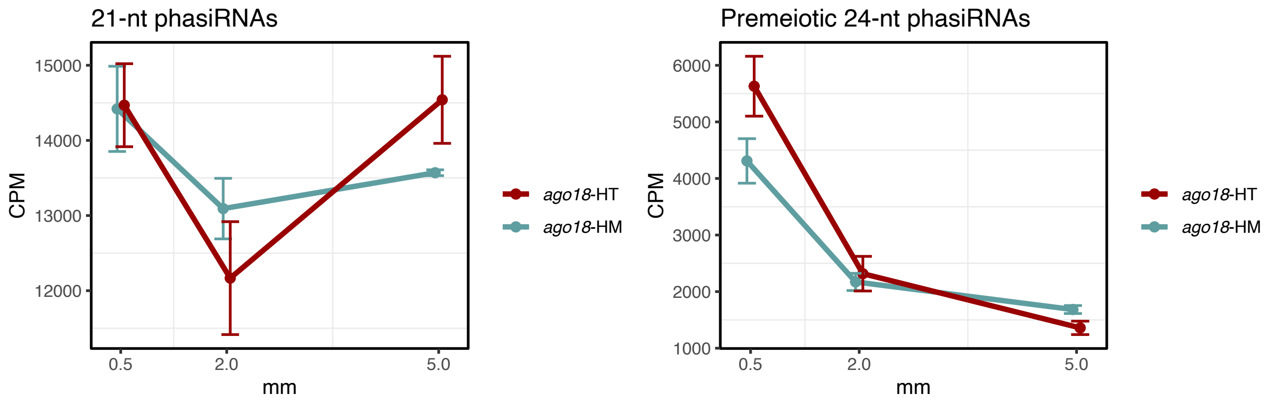


**B**

| | 0.5 mm | 2 mm | 5 mm |
|---|---|---|---|
| 21-nt phasiRNA | 0.953979698 | 0.355797547 | 0.23542009 |
| Premeiotic 24-nt phasiRNA | 0.121698813 | 0.700015424 | 0.093317272 |
| Meiotic 24-nt phasiRNA | 0.495300153 | 0.343272602 | **0.003812074** |

### Figure S13

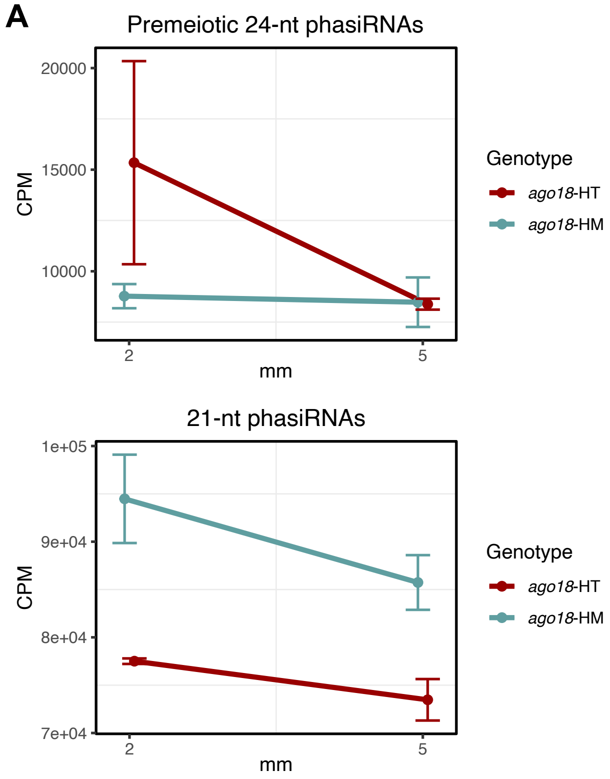


**B**

| | 2 mm | 5 mm |
|---|---|---|
| Premeiotic 24-nt phasiRNA | 0.318801764 | 0.945713879 |
| Meiotic 24-nt phasiRNA | **0.045561126** | **0.003091776** |
| 21-nt phasiRNA | 0.167977163 | **0.02996213** |

### Figure S14
